# Evolutionary trends in the emergence of skeletal cell types

**DOI:** 10.1101/2024.09.26.615131

**Authors:** Amor Damatac, Sara Koska, Kristian K Ullrich, Tomislav Domazet-Lošo, Alexander Klimovich, Markéta Kaucká

**Affiliations:** Max Planck Institute for Evolutionary Biology, August-Thienemann-Straße 2, 24306 Plön, Germany; Christian Albrechts University Kiel, Am Botanischen Garten 1-7, 24118 Kiel, Germany; Laboratory of Evolutionary Genetics, Division of Molecular Biology, Ruđer Bošković Institute, Bijenička cesta 54, HR-10000 Zagreb, Croatia; School of Medicine, Catholic University of Croatia, Ilica 242, HR-10000 Zagreb, Croatia

**Keywords:** cell type evolution, single-cell phylotranscriptomics, skeletogenesis, hypertrophic chondrocytes, lineage-specific genes

## Abstract

The emergence of novel cell types fuels evolutionary innovations and contributes to the diversity of life forms and their morphological and functional traits. Cell types are fundamental functional units of multicellular organisms defined by their specific gene expression programs. The evolution of these transcriptional programs is driven by genetic changes, such as gene co-option and cis- regulatory evolution, known to facilitate the assembly or rewiring of molecular networks and give rise to new cell types with specialized functions. However, the role of novel genes in this complex evolutionary process is underexplored. Here, we examine the trends in skeletal cell type evolution with a focus on lineage-specific genes. We find that immature chondrocytes express the oldest transcriptome and resemble ancestral skeletogenic cell type, supporting the existence of a conserved genetic program for cartilage development in bilaterians. The subsequent acquisition of lineage-restricted genes led to the individuation of the ancient gene expression program and powered the emergence of osteoblasts and hypertrophic chondrocytes. We found a significant enrichment of Vertebrate-specific genes in osteoblasts and Gnathostome-specific genes in hypertrophic chondrocytes. By identifying the functional properties of the recruited genes, coupled with the recently discovered fossil evidence, our findings challenge the long-standing view on the evolution of vertebrate skeletal structures and suggest that endochondral ossification and chondrocyte hypertrophy evolved already in the last common ancestors of gnathostomes. Finally, our findings highlight the critical role of novel genes in shaping cellular diversity.

## Introduction

During the evolutionary history of animals, a series of major evolutionary transitions were facilitated by the emergence of novel traits, which led to the enormous diversity and complexity of extant animals. The emergence of new cell types and their further individuation contributed to the origin of such functional and morphological novelties[1–3]. Cell types are considered functional and evolutionary units that fuel innovations along the phylogeny of animals[4, 5]. From an evolutionary standpoint, a cell type is a group of cells that share morphological and functional features and possess specific gene expression profiles that have arisen through the evolution of a particular regulatory program[6]. Such regulatory programs are assembled through the integration of existing molecular modules, incorporation of novel genes, and evolution of the cis-regulatory landscape[6–8]. These mechanisms jointly contribute to the emergence of novel cell types and their further radiation. The cell types that evolved from the same ancestor usually share the expression of many genes[9, 10]; therefore, the knowledge of the evolutionary origin of each gene is instrumental in understanding the origin, homology, and functional properties of a cell type.

The evolution of cell types has been conventionally studied by comparing the expression of a relatively small number of mostly conserved marker genes representing a cell type identity[11–14]. These investigations generated fundamental knowledge of cell type evolution and proposed homology between numerous cell types. However, assessing such a limited set of genes does not provide a comprehensive overview of cell type evolution[9] since the same gene can be a marker of several unrelated cell types. To complement the marker gene approach, alternative strategies that identified the core sets of co-regulated genes and used them as molecular signatures of certain cell types were employed[15]. The recent emergence of single-cell phylotranscriptomics, in particular combining single-cell transcriptomics and genomic phylostratigraphy, offers a new avenue to explore principles of cell type evolution by investigating the link between gene and cell type origin[16, 17]. A cell expresses thousands of genes, and comprehensively analyzing the evolution of its entire gene expression profile can provide even more detailed insights into the cell type’s origin and history.

The evolution of skeletal cell types had a major impact on the evolutionary novelties, adaptations and transitions across animal lineages. In this study, we employed the skeletogenic lineage of developing murine hindlimb as a model because it comprises diverse cell types that generate stiff elements in the process of endochondral ossification. In contrast to the formation of elastic and fibrous cartilage or dermal bones, where only few cell types are involved, endochondral ossification is a bone-forming mechanism that employs the majority of skeletal cell types known in vertebrates, both from the chondrogenic and osteogenic lineages (Fig. 1a). In vertebrates, chondrocytes produce cartilage that forms an embryonic blueprint of the future skeleton[18]. In most vertebrates, with the exception of cartilaginous fishes (Chondrichthyes), this template is subsequently substituted by mineralized bone. Chondrocytes that form cartilaginous supportive elements are also present in diverse invertebrate lineages (Fig. 1a)[2, 19, 20], as was deducted from the shared expression of several marker genes and biochemical properties[14, 20]. However, the discontinuous presence of cartilage along the invertebrate phylogeny raised a question about the evolutionary origin of this program (Fig. 1a)[14, 21].

**Fig. 1.**
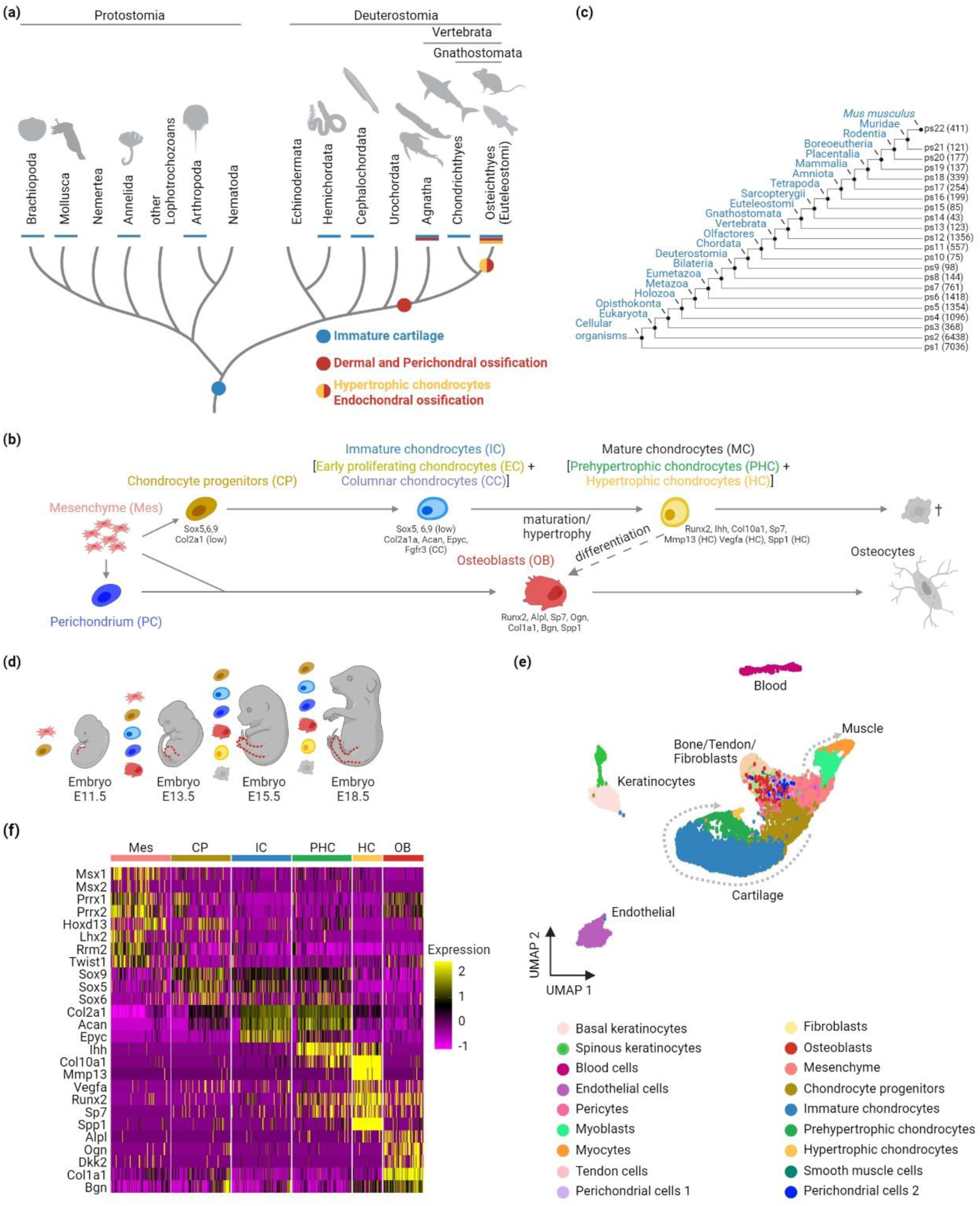
Paradigm and methodology to explore the contribution of novel taxon-restricted genes to the evolution of skeletogenic cell types. a) The evolution of skeletal cell types in bilaterians. Colored dots indicate the current view on the evolutionary origin of cartilage (blue), dermal, perichondral and endochondral bone (red). Chondrocyte hypertrophy (orange) is observed in Euteleostomi. Colored bars indicate the presence of skeletal tissues in a given lineage. b) Schematic representation of cell types and their differentiation trajectories involved in the endochondral ossification during embryonic development. c) Phylostratigraphy map of the mouse protein-coding genes. Each phylostratum (ps) number corresponds to a node in the phylogeny. Numbers in the parentheses indicate the number of genes originating in the phylostratum. d) Embryonic stages covered by single-cell transcriptome dataset from developing mouse hindlimb. Stage E11.5 contains mostly mesenchymal condensations; E13.5, E15.5, and E18.5 comprise cell types typically involved in chondrogenesis and osteogenesis in the limb. Mesenchymal cells condense within the developing limb bud and differentiate into chondrogenic and osteogenic cells. In the maturation process, chondrocytes undergo programmed cell death or contribute to the osteogenic lineage through transdifferentiation. e) Single-cell transcriptome analysis. The dataset was reanalyzed using the Seurat pipeline, and the cell clusters were projected using two-dimensional UMAP. Individual colors indicate the cell cluster identities. f) Gene expression profile of skeletogenic cell clusters using cell type-specific genes.

In embryonic development, both chondrocytes and osteoblasts originate primarily from mesenchymal cells, whereby the chondrogenic lineage exhibits more differentiation complexity (Fig. 1b). The chondrocyte differentiation trajectory spans from mesenchymal progenitors through proliferating immature chondrocytes (IC), to terminally differentiated mature chondrocytes (MC)[22, 23]. IC can be further divided into early chondrocytes (EC, also known as round or resting chondrocytes) and columnar chondrocytes (CC, also referred to as proliferating or flattened chondrocytes) while MC comprise prehypertrophic (PHC) and hypertrophic chondrocytes (HC) (Fig. 1b)[18, 24]. Each of these cell types is characterized by a specific combination of marker genes (Fig. 1b). The unique property of maturing chondrocytes in endochondral ossification is their ability to undergo a programmed cell death defined by distinct morphological and transcriptomic changes[25, 26], referred here to as the hypertrophy. An alternative cell fate trajectory of MC is their differentiation into osteoblasts (OB) (Fig. 1b)[27–29].

While vertebrate IC are considered homologous to those found in nonvertebrate lineages[30], MC and endochondral ossification are only observed in extant bony vertebrates – Euteleostomi/Osteichthyes[20, 30–33]. Osteoblasts that directly differentiate from the mesenchymal cells first appeared already in stem vertebrates and produce dermal and perichondral bones (Fig. 1a)[32–35]. This renders endochondral ossification an evolutionary younger process compared to dermal and perichondral ossification (Fig. 1a). Chondrocytes and osteoblasts are known to share the expression of many genes[36–38], suggesting that these cell types originate from a common evolutionary ancestor. It has also been hypothesized that osteoblast’s transcriptome evolved from the chondrogenic gene expression program; however, compelling evidence is missing, and the evolutionary dynamics of their transcriptome assembly and individuation are yet to be deciphered[24, 30, 39].

Here, using the recently emerged phylotranscriptomics, integrating genomic phylostratigraphy and single-cell transcriptomics, we strive to deepen the knowledge of skeletal cell type evolution. In particular, we ask how the evolution of lineage-restricted genes contributed to the emergence of novel cell types and their biological function.

## Results

To estimate the evolutionary age of genes, we employed genomic phylostratigraphy, also known as “gene birthdating”, that determines the evolutionary origin of every protein-coding gene from a focal species by tracing significant shifts in the protein sequence across the tree of life[40, 41]. Genomic phylostratigraphy assigns a relative age to each gene according to the most distant phylogenetic node where sequence homology is still detectable[42]. We carried out a phylostratigraphic analysis of all mouse protein-coding genes across 22 phylogenetic nodes (phylostrata, ps), starting from the common ancestor of cellular organisms to the terminal branch, *M. musculus* (Fig. 1c; Supplementary Fig.1; Supplementary Tables1-3). The generated phylostratigraphic map indicates the evolutionary origin of murine 22,590 genes along the consensus phylogeny, and the obtained distribution is similar to the earlier observations[43, 44]. Previous studies utilized genomic phylostratigraphy to explore the evolution of developmental stages, organs, tissues and cells acquired from heterogeneous samples that contained a mixture of different cell types[45–50], demonstrating its capability to provide evolutionary insights into developmental processes. Therefore, combining information on gene origin with transcriptome data at a single cell level provides an opportunity to better understand the evolution of cell type-specific transcriptomes. Here, we applied genomic phylostratigraphy in the context of single-cell transcriptomics of the skeletal cell types.

To capture the gene expression program of skeletogenic cell types, we took advantage of a single-cell transcriptome dataset from murine hindlimb development (Fig. 1d)[51]. With in-depth analysis and annotation of cell clusters (Fig. 1e-f; Supplementary Fig. 2), we identified all relevant cell types, including IC, PHC, HC, and OB. The remaining clusters represent endothelial, myogenic, and hematopoietic lineages (Fig. 1e; Supplementary Fig. 2). We emphasize that the great advantage of the single-cell transcriptomes, compared to the bulk sequencing data, is that each cell type is represented by dozens or hundreds of individual cells, which serve as independent biological replicates. This allows statistical comparison among cell types and uncovers variation in evolutionary signals between individual cells, which is inherently invisible in pooled samples[17]. The existing single-cell transcriptomic technologies do not detect all genes within the cell’s transcriptome; however, they provide the most comprehensive information on the composition of individual transcriptomes.

To estimate the relative evolutionary age of the cell type-specific transcriptomes, we calculated the transcriptome age index (TAI)[45] for each skeletogenic cell. TAI integrates the average age of all protein-coding genes expressed in a cell with their expression level. A lower TAI value indicates an evolutionarily older transcriptome, while a higher TAI value corresponds to an evolutionarily younger transcriptome. We compared the TAI values of all cells among skeletogenic cell types (Fig. 2a; Supplementary Fig.3).

**Fig. 2.**
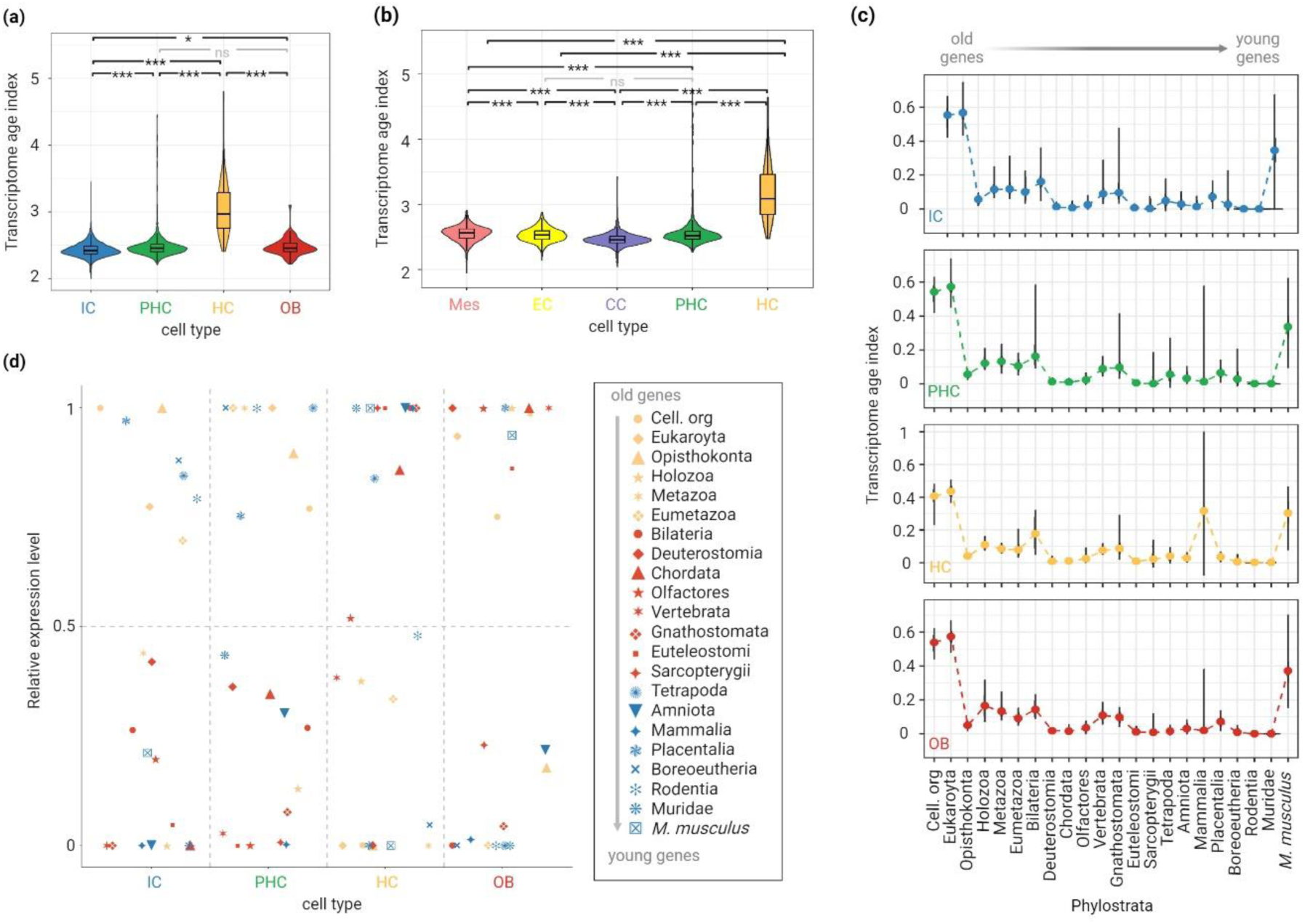
Phylotranscriptomic analysis of skeletal cell types. a) Phylogenetic age of the skeletal cell types based on the transcriptome age index (TAI). A lower TAI value indicates a phylogenetically older transcriptome. All expressed genes in a given cell type are considered. Of note, the TAI calculation for OB was done irrespective of their cellular origin (whether OB differentiated from perichondrium, mesenchymal cells or HC is not distinguished) b) Phylogenetic age of the cell types of chondrogenic lineage, clustered in higher resolution to distinguish individual differentiation stages, based on the transcriptome age index (TAI). c) Partial TAI (pTAI) of skeletal cell types split according to the origin of the genes from the different phylostrata. The violin plot in individual phylostrata shows the TAI values for every cell, while the line plot connects the median value in each phylostrata. d) Relative expression of genes from different phylostrata across skeletal cell types. The mean expression of genes for a given ps in each cell type was linearly transformed to a relative expression interval (minimum expression= 0, maximum expression= 1). a,b) The statistical significance of differences between TAI values was evaluated using a pairwise Wilcoxon test corrected for multiple comparisons by Benjamini and Hochberg (BH). Asterisks denote adjusted p-value levels (* ≤0.05, **≤0.01, ***≤0.001). IC, Immature chondrocytes; PHC, Prehypertrophic chondrocytes; HC, Hypertrophic chondrocytes; OB, Osteoblasts; EC, Early chondrocytes; CC, columnar chondrocytes.

First, we observe that chondrocytes (comprising both IC and MC) express, on average, older transcriptomes than OB (Supplementary Fig.3a). This is particularly evident when only the TAI of upregulated differentially expressed genes are compared between these cell types (Supplementary Fig.3a). This result agrees with the molecular, phylogenetic, and fossil evidence that cartilage is evolutionarily older than bone[14, 20, 21, 52, 53].

Because the chondrocyte population is heterogeneous and comprises both mature and immature states, we analyzed this population at a higher resolution and clustered the cells according to their maturity (ontogeny). The TAI of immature chondrocytes (IC) is lower than the TAI of mature chondrocytes (comprising PHC and HC) and osteoblasts (OB) (Fig. 2a; Supplementary Fig.3b). Interestingly, HC demonstrates substantially higher TAI values than OB and all other chondrocyte cell types. These patterns remain consistent when the number of cells for each cell type was randomly downsampled (n=50, n=100) (Supplementary Fig.3c-d), indicating the TAI is not biased due to the unequal number of cells among cell types. As the TAI calculation incorporates gene expression level as a weight factor[17, 45], these results indicate that the higher TAI of OB and HC are influenced by higher expression of recently evolved genes as compared to IC. Most notably, our findings suggest HC has the youngest transcriptome among the skeletal cell types, even younger than OB. By partitioning the immature chondrocyte population even further, we observe that CC, representing the last immature chondrogenic state, express a transcriptome that appears older than any tested cell type (Fig. 2b). This suggests that CC transcriptome might resemble the oldest chondrogenic (or skeletogenic) genetic program among the skeletal cell types.

To reveal the contribution of the individual phylostrata to the TAI value, we calculated the partial TAI (pTAI) scores according to the origin of genes from the different phylostrata[45] (Fig. 2c; Supplementary Fig.4). To simplify the presentation of these results, we connected the median of partial TAI values in each phylostrata for each skeletal cell type (Fig. 2c). We observe that genes originating from cellular organisms (ps1) to Bilateria (ps7) contribute greatly to the global TAI values. Notably, these ancient genes include *Sox*, *Runx* and collagen gene families that are conventionally used as marker genes for skeletal cell types (Supplementary Table1)[39, 54]. This highlights that the marker genes of skeletal cell types consist of ancient genes that have been recruited into their transcriptomes during evolution. Consequently, this observation supports that the ancient core genes involved in the first skeletal (chondrogenic) gene expression program were assembled during the emergence of Bilateria and have been conserved, as evidenced by the existence of cartilage in several protostome and deuterostome lineages[14, 20]. Comparison of pTAI values among skeletal cell types (Supplementary Fig.4) reveals that most ancient genes (cellular organisms-Opisthokonta) are more expressed by IC, further supporting that IC is reminiscent of the ancient chondrogenic genetic program.

We find that genes that have emerged later in the evolution, including genes from Vertebrata (ps11), Gnathostomata (ps12), and *M. musculus* (ps22), contribute greatly to the global TAI score of OB and HC (Fig. 2c). Particularly, OB show a significantly high pTAI at the onset of Vertebrata (ps11), while HC exhibit considerably elevated pTAI in Gnathostomata (ps12) (Supplementary Fig.4). These observations suggest that the corresponding phylogenetic nodes were important periods in the evolution of OB and HC, respectively, and contributing to the transcriptome divergence from the ancestral skeletogenic genetic program. We also detect a strong surge of pTAI in HC at the origin of Mammalia (ps17), where solely a high expression of *Spp1* (Osteopontin), a marker of HC, causes the substantial pTAI increase. This indicates that *Spp1* is an important component of the HC gene expression program, and its high expression is required for mineralization and bone properties[55].

To better understand the phenomenon of ancestral cell type’s transcriptome divergence, we compared the proportions of expressed genes shared by IC, HC, and OB along the ps nodes or unique to each cell type (Supplementary Fig.5). We observe that IC share a large proportion of genes with both HC and OB. However, HC and OB share fewer genes. To comprehend the functional evolutionary relationships among these cell types, we performed a Gene ontology (GO) enrichment analysis on the unique and shared genes among IC, OB, and HC (Supplementary Fig. 6; Tables 4-9). We find that genes unique to IC are associated with the known properties of cartilage, such as cartilage development and synthesis of proteoglycan and aminoglycan. HC and IC share genes associated with cartilage development, while OB and IC share genes related to ECM organization. On the other hand, unique genes for OB and HC are associated with their biological properties of becoming a bone and mature cartilage, respectively. These include ECM organization, ossification, limb and skeletal development for OB, and cell death, ossification, cartilage and bone development, cytoskeleton organization, fat cell differentiation and mesenchyme development for HC. Given that IC’s transcriptome likely resembles the ancestral cell type gene expression program, these findings suggest that HC and OB transcriptomes evolved independently from a common ancestral (chondrogenic, IC-like) gene expression program. Coherently, functions of genes shared between OB and HC are linked to their role in endochondral ossification, an evolutionary younger ossification mechanism that is observed among extant Osteichthyes[32]. Overall, our observations suggest the further elaboration of the ancestral skeletogenic gene expression toolkit, whereby the recruitment of lineage-specific genes may facilitate transcriptome individualization.

To further demonstrate to what extent genes of particular evolutionary origin contribute to the overall cell type-specific transcriptomic profile, we performed a relative expression analysis of genes evolved in different phylostrata (Fig. 2d). In contrast to TAI analysis, relative expression analysis does not apply different weight on each phylostratum but only considers the expression of genes in every phylostratum. We show that IC and PHC highly express mainly ancient genes that originated between the origin of cellular organisms and the radiation of Eumetazoa (ps1-6, orange shapes). In contrast, evolutionarily younger genes that emerged in Bilateria (ps7) or later (red and blue shapes) are more expressed by either OB or HC. Comparisons of the average expression of genes from each ps and among cell types provide statistical support that older genes tend to be more expressed by IC, while younger genes are more expressed in OB and HC (Supplementary Fig.7). Taken together, the analysis of shared and unique gene composition (Supplementary Fig.5-6) and relative expression of the genes originating in different phylostrata (Fig. 2d; Supplementary Fig.7) highlight the contribution of evolutionarily younger genes to individualization of OB and HC gene expression programs and support their more recent evolutionary origin compared to IC.

TAI is a measure that considers all gene expression levels via their partial concentrations[45]. However, transcriptome data could also be assessed by looking for statistical enrichment of all upregulated genes from each phylostratum regardless of the gene expression levels in a cell. This type of analysis gives equal weight to highly expressed genes and those represented by a small number of transcripts[56]. We thus performed enrichment analysis on all upregulated genes of skeletal cell types transcriptomes (Fig. 3a). We observe significant enrichments in several phylostrata such as Holozoa (ps4; for both OB and HC), Metazoa (ps5; for HC), Bilateria (ps7; for OB), Mammalia (ps17; for OB) and Placentalia (ps18; for IC). Interestingly, a significant number of recently evolved genes that are part of the OB and HC transcriptomes appeared at the onset of Vertebrata (ps11) and Gnathostomata (ps12), respectively, which aligns well with patterns previously deducted from TAI and relative expression analysis. These results further show that novel genes originating in ps11 and ps12 were recruited specifically into the OB and HC gene expression programs. While our analysis cannot demonstrate whether these genes were incorporated into the gene expression programs at the time of the genes’ origin, it indicates the earliest point when such integration was possible. Our findings suggest that the genes from the enriched phylostrata, particularly novel genes, are involved in the adaptive history of skeletal cell types. For OB, enrichment in ps11 strongly coincides with the reported first emergence of bones in Vertebrates[32, 35]. In the case of HC, the significant enrichment of genes in ps12 suggests that HC originated already in the ancestor of jawed vertebrates (Gnathostomata) instead of bony vertebrates (Osteichthyes) [57]. Unfortunately, the species from the base of stem gnathostomes are extinct, which prevents us from carrying out direct validation. Considering the indispensable role of HC in endochondral ossification, the recent fossil discovery of endochondral bone in a placoderm-like fish, Minjinia turgenensis[58], provides strong support to our finding.

**Fig. 3.**
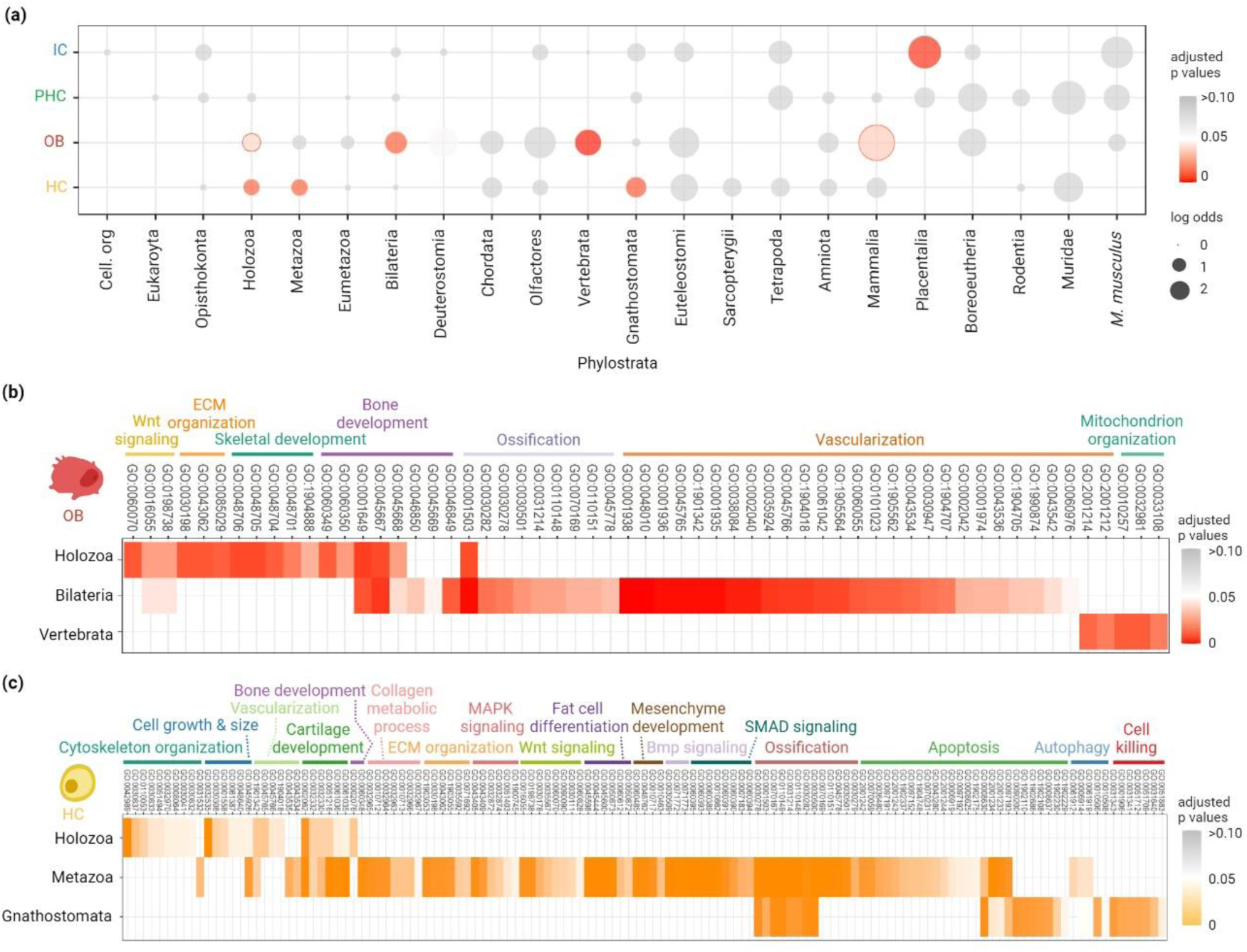
Functional enrichment of genes associated with the evolution of skeletal cell types. a) Enrichment analyses on upregulated DEG distribution across the phylogeny. The representation of novel genes associated with the evolution of a particular cell type is shown for each ps in log-odds values. Red circles indicate statistically significant enrichment of genes associated with the evolution of a particular cell type, while grey circles indicate enrichments that are not statistically significant. Enrichments were tested using a two-tailed hypergeometric test corrected for multiple comparisons by FDR (p ≤ 0.05). b,c) GO enrichment analysis for biological processes of DEGs originated from the significantly enriched phylostrata in Fig. 3a. All enrichment analyses were corrected by BH (p ≤ 0.05). Enriched phylostrata, including Placentalia (IC) and Mammalia (OB), did not provide significant GO terms. A complete list of statistically significant GO terms is provided in the supplementary tables.

To better comprehend the molecular and functional modules associated with the evolution of OB and HC, we perform GO enrichment analysis on the upregulated genes from each enriched phylostrata (Fig. 3b-c; Supplementary Tables 10-15). For OB, the genes originating in older phylostrata (Holozoa and Bilateria) are associated with Wnt signalling, extracellular matrix (ECM) organization, bone development, ossification, and vasculogenesis. These include known OB transcription factors from *Runx* and *Twist* gene families arising in Holozoa and ossification- and vasculogenesis-related genes such as vascular endothelial growth factor (*Vegf*) originating from Bilateria. OB furthermore utilize genes related to mitochondrion organization that emerged in Vertebrata (ps11), which may be required for the specific energetic and metabolic demand during the mineralisation process [58]. Notably, OB additionally employ novel vasculogenesis-related genes *Ramp2* and *Tmem100* from the node Vertebrata (ps11) that enabled the recruitment of the pre-existing vasculogenesis module into the new context of bone formation[59, 60]. Vascularization, a hallmark feature of bone formation, is considered a major evolutionary innovation that distinguishes bone from the ancestral skeletal tissue, the non-vascularized cartilage[61, 62]. Overall, these newly evolved components of transcriptional modules and the associated functions align well with the described biology of bone formation.

In Holozoa (ps4), HC-associated genes were assigned to GO terms linked with cytoskeleton organization, cell growth and size, which may be related to the ability of chondrogenic cells to change their shape and increase in size – a hallmark feature of HC during chondrocyte maturation process[63, 64]. The Holozoan genes are additionally associated with GO terms related to cartilage and bone development. Noteworthy among them are transcription factors from the *Runx* and *Smad* family, recognized for their essential roles in the development of mature cartilage and bone [65–67]. HC-associated genes of Metazoan origin (ps5) are linked to key signalling pathways such as *Wnt*, MAPK, Bmp and SMAD that are known to facilitate chondrocyte differentiation and maturation[68]. HC also utilize holozoan and metazoan genes related to bone development and ossification, which align well with the HC’s involvement in endochondral ossification. Ancient metazoan genes that are part of the HC transcriptome, including *Bax, Cflar, Mcl1, Lgals3, Wfs1, Pabpn1, Nr4a2, Wnt4 and Ackr3*, are linked to the apoptotic pathway. Notably, the HC transcriptome is further complemented by genes of Gnathostomata (ps12) origin that are known to modulate programmed cell death, such as *Bid, Bcl2l1, Bmf, Ier3, and Muc1*. These observations provide the first molecular evidence that the evolution of HC was fueled by the acquisition of cell death control, a hallmark process of HC during hypertrophy[25, 28]. In summary, our findings indicate that the evolution of skeletal cell types (OB and HC) followed a general principle of transcription factor recruitment, together with the co-option of ancient modules (vasculogenesis and cell death) and the incorporation of novel genes. The analyses pinpointed novel vertebrate- and gnathostome-specific genes that equipped evolutionarily younger skeletal cell types, OB and HC, with specialized functions.

To test whether we can recapitulate the evolutionary trends deciphered using the mouse limb development paradigm are generally valid for Vertebrata, we complemented our study using a zebrafish (*Danio rerio*) model. This model represents a particular interest due to its phylogenetic position as a species most removed from the mouse within the Osteichthyes, for which a single-cell dataset comprising a spectrum of skeletogenic cell types is currently available. Notably, the zebrafish skeleton is mainly generated by intramembranous ossification; therefore, capturing HC, in particular, proved challenging. In short, we replicated the phylotranscriptomic analysis using two datasets of fish development resolved at the single cell level, profiled the TAI scores in diverse skeletogenic cell types, and performed enrichment analysis (Supplementary Result, Supplementary Fig. 16-18). Overall, the results gained from phylotranscriptomic analysis in zebrafish corroborate our main findings in mouse. Most remarkably, the gene ontology analysis again pointed to the enrichment of ECM and cell oxygen supply categories in OB, as well as ECM regulation and cell death control in the HC. In sum, our findings on the zebrafish dataset support the concept of step-wise recruitment of novel lineage-specific genes contributing to the emergence of cell types.

## Discussion

Cell type emergence is a complex evolutionary process simultaneously powered through co-option, novel gene acquisition, and the evolution of regulatory non-coding elements[7, 69–75]. Here, we set out to understand the contribution of different phylogenetic levels via protein-coding genes to the evolution of skeletogenic cell types. The evolution of skeletal tissues has been studied for decades, allowing us to compare our phylotranscriptomic findings with the extensive evidence from paleontology, phylogenetics, and molecular research. The current knowledge of skeletal cell type evolution is mainly deducted from the comparative biochemical, histological and marker gene expression surveys. Previous investigations have assessed the expression of a limited number of selected marker genes from *Sox, Runt*, hedgehog, and collagen gene families to deduce the cell type origin[13, 14, 53, 76–78]. However, these gene families are evolutionarily ancient and might be expressed by multiple unrelated cell types[79–81]. While these studies uncovered numerous commonalities among skeletal tissues in both vertebrate and invertebrate lineages (Fig. 1a)[14, 21], they raised questions about mechanisms fueling the skeletal cell type evolution, whether the skeletal cell types evolved from a common ancestral cell type or how their genetic programs diverged. Here, we demonstrate the potential of single-cell phylotranscriptomics in addressing such inquiries, focusing particularly on how the evolution of lineage-restricted genes contributed to the emergence of novel cell types and their functional properties. We explored the evolutionary history of the gene expression programs operating in skeletal cell types and pinpointed the essential role of novel genes in this evolutionary process.

We exemplify how the TAI, relative expression, enrichment, and GO term analyses offer deeper insights into the evolution of skeletal cell types transcriptomes. The transcriptome age index assumes that cell types expressing older transcriptomes (having lower TAI) likely retained the genetic program from ancestral cell types and tend to be functionally conserved, while cell types expressing young transcriptomes utilize novel genes and evolve later[17]. Therefore, IC expressing the oldest transcriptome likely represents the prototypical skeletal (chondrogenic) expression program conserved among bilaterians. This finding is corroborated by the similarities of the immature cartilage across protostome and deuterostome lineages in the structural and biochemical properties and expression of core marker genes[14]. We conclude that OB and HC likely emerged later in the phylogeny through an independent divergence and individuation from the ancestral gene expression program (Fig. 4). HC expresses a younger transcriptome compared to OB, which supports the observed distribution of bones and mature cartilage on the vertebrate phylogeny (Fig. 1a). However, the HC gene expression program appears to be assembled earlier than previously thought. Our findings indicate that the last common ancestor of Gnathostomes likely possessed a virtually complete HC gene expression toolkit (Fig. 4a, c). Since chondrocyte hypertrophy is exclusively involved in endochondral ossification, it is evident that HC and endochondral ossification emerged simultaneously. Our results, strongly supported by the recent fossil evidence of endochondral bone in a stem Gnathostome[82], render HC and endochondral ossification as Gnathostome-specific novelty (Fig. 4c). Consequently, these discoveries support the emerging assertions that Chondrichthyes lost the ability to make bones [82, 83]. Although the interpretation of TAI could be subject to debate and may not be universally applicable to all cell types, in this study, the combination of TAI, relative expression, and particularly enrichment analyses, supported by published knowledge, jointly provide insights into the evolutionary origin of skeletal cell types.

**Fig. 4.**
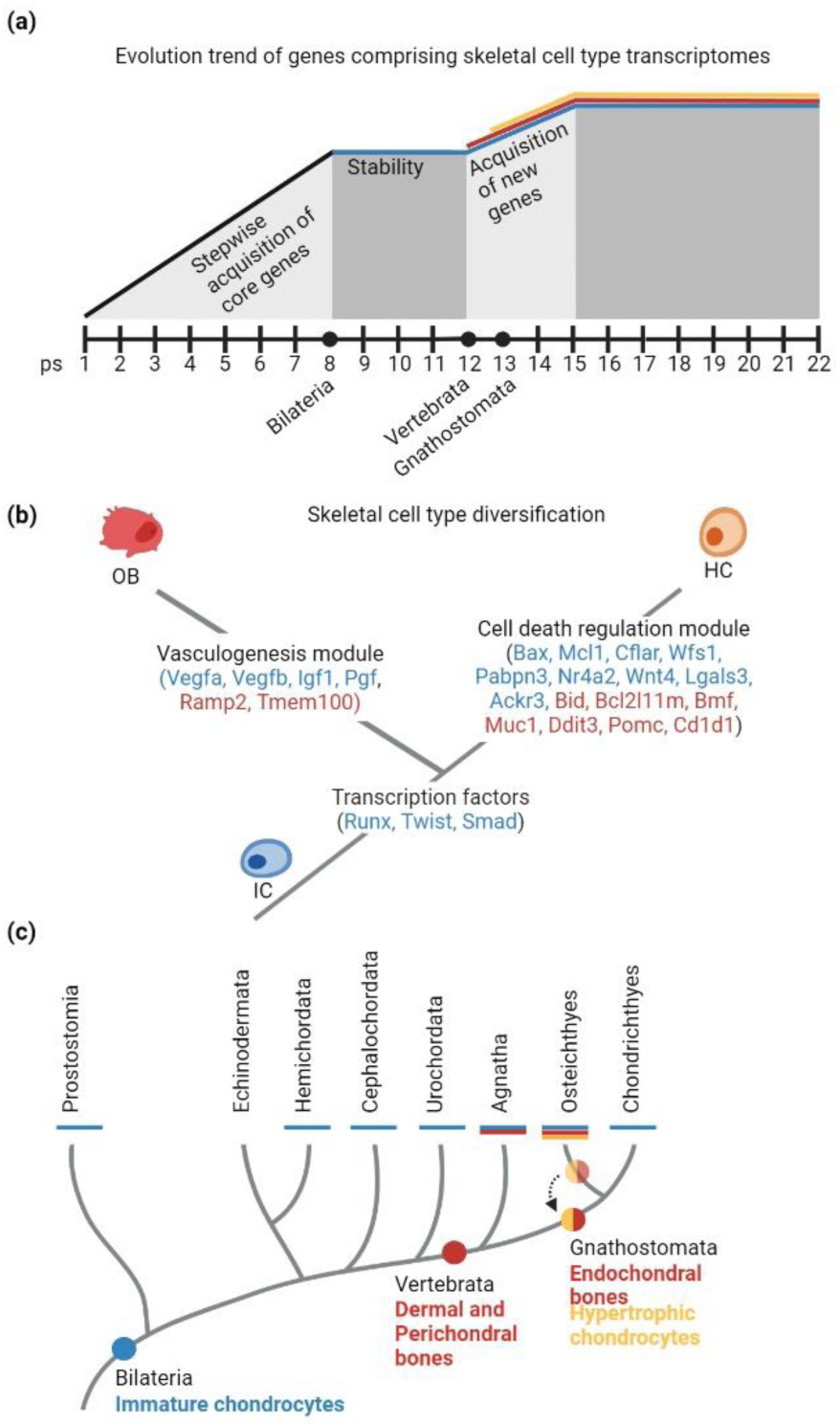
The molecular evolution and origin of skeletal cell types. a) Model and order of the emergence of skeletal cell type-specific gene expression programs. The ancestral proto-skeletal (chondrogenic) program emerged at the onset of Bilateria after the assembly of ancient functional modules that appeared during the origin of cellular organisms to the ancestors of bilaterians. Skeletal cell types diversified at the onset of vertebrates after divergence events from the ancestral skeletal program. Y-axis is not scaled against geological time. b) Skeletal cell type diversification occurred through the recruitment of transcription factors and the integration of additional functional modules. We found that the implementation of genes modulating vasculogenesis and mitochondrion organization in OB and programmed cell death in HC contributed to the individuation of the ancestral IC expression program. c) Simplified phylogenetic origin of skeletal cell types in animals. We propose that endochondral ossification and HC evolved in the common ancestor of Chondrichthyes and Osteichthyes, moving the origin of HC from the ancestors of bony vertebrates.

Examining the entire cell type-specific transcriptomes allowed us to shed light on their evolutionary step-wise assembly and uncover the contribution of the ancient and novel genes to the emergence and individualization of skeletal cell types’ gene expression programs. Here, we identify genes originating from particular evolutionary nodes that provided the cell types with new distinct biological identities and functions. This facilitates the individuation from the ancestral chondrogenic genetic program and drives the emergence of evolutionarily younger cell types, OB and HC. We show that the recruitment of transcription factors from Holozoa (*i.e.*, *Runx, Twist, Smad*) contributed to the cell type diversification from the ancestral chondrogenic cell type (Fig. 4b), supporting the proposed principle that the evolution of cell types requires changes in transcription factors to create new cell type identity. In conjunction, the employment of novel protein-coding genes enables the integration and modulation of ancient processes (pre-existing biological functions) as new cellular modules of HC and OB (Fig. 4b). The recruitment of genes controlling the ancient programmed cell death machinery facilitated the evolution of HC, while genes modulating vasculogenesis propelled the emergence of OB. While the functional role of individual genes is yet to be experimentally validated, these findings support the notion that cell type evolution is linked to the acquisition of new functions[84]. Interestingly, the origin of lineage-restricted genes comprising the specific cell type transcriptome coincides with the emergence of their respective cell types (Vertebrate-specific genes in OB; Gnathostome-specific genes in HC), providing a mechanistic link between the evolution of novel genes, acquisition of new cellular functions and cell type emergence. While the applied research strategy cannot reveal whether the novel genes were incorporated into the cell type-specific gene expression program at the time of the genes’ origin, it pinpoints the very first moment when such integration was possible. However, the presented finding is supported by a prior study on a considerably smaller phylogenetic clade of Cnidaria, where the authors demonstrated that the recruitment of novel genes fuels the emergence of a new cell type[16]. We acknowledge that extensive sampling of skeletogenic tissues across phylogeny will be required to verify the evolutionary assembly of skeletal cell type transcriptomes. However, single-cell transcriptome datasets from non-model species across the tree of life are still relatively rare and do not contain skeletal cells. In the future, comparative single-cell phylotranscriptomics of corresponding skeletal tissues across the phylogeny will clarify the time of the genes’ integration.

The single-cell phylotranscriptomics of skeletal cell lineages in a mouse model enabled us to formulate a theory on skeletal cell type evolution. However, the study is currently limited by using data only from developing murine hindlimbs. To acquire a thorough understanding of how the entire diversity of skeletal cell types evolved, our findings must be corroborated 1) through interspecies comparisons and 2) from all skeletal tissues (containing mature osteocytes, osteocytes of dermal and perichondrial origin, chondrocytes forming hyaline, elastic and fibrocartilage). At the time of this study, data from other species comprising IC, OB and HC were unavailable. Therefore, we acknowledge the importance of taxonomic sampling in drawing more solid interpretations in the future. Nevertheless, our main conclusions on the evolutionary origin of skeletal cell types align seamlessly with published knowledge and new fossil evidence.

In summary, we present a study that may inspire the design principles for future investigations tailored specifically to questions of a particular cell type evolution and individuation of gene expression programs from their ancestor. The recent advances in single-cell omics technologies deliver complex and high-resolution information on the molecular profile of individual cells. Examining several thousands of genes in each cell provides a comprehensive insight into the cell identity along a developmental trajectory[85]. The single-cell phylotranscriptomics represents a universal approach that can be applied to different models and paradigms. The implementation of additional tools for inferring gene homologies, such as BLAST- independent HMM approaches (for instance, see [86–88]), incorporation of an ever-increasing knowledge on novel gene emergence, and methodologies to distinguish between duplication-divergence and de novo evolution, the synteny analysis, will promote a broader and even more confident application of genomic phylostratigraphy on single-cell data to address questions of cell type evolution. This will be particularly important for studies of cell type evolution in species with duplicated genomes, where the contribution of individual gene paralogues needs to be considered.

## Methods

### Single-cell transcriptome dataset

We employed published 10x Genomics single-cell transcriptome datasets from mouse and zebrafish. In the mouse dataset, samples of the developing hindlimb were recovered from four embryonic times (E11.5, E13.5, E15.5, and E18.5)[51]. Additionally, we utilized two zebrafish datasets: one covering the larval stages 5dpf and 14dpf and comprising cell progenies that emerged from labeled neural crest cells in the head region[89], and the second dataset analyzing larval stage 5 dpf and a whole embryo[90].

### Single-cell transcriptome analysis

In the single-cell dataset of mouse limb development, the raw count matrix was processed using the Seurat v3[91] R package. We filtered out cells with less than 200 and more than 5,000 genes. Cells with more than 20% mitochondrial read content were also removed. The genes with either zero expression in all cells or only expressed in less than ten cells were excluded. The filtered data were normalized and variance stabilized (cell cycle phase and mitochondrial percentage regression) using the SCTransform algorithm[92]. The stages were then integrated using the highly variable genes identified using SCTransform. To visualize the integrated data, the resulting file was subjected to dimensionality reduction methods such as Principal Component Analysis (PCA) and Uniform Manifold Approximation and Projection (UMAP). We clustered the cells using Seurat’s default settings (K-nearest neighbour (KNN) graph and Louvain algorithms). The cell clusters were identified using the upregulated differentially expressed genes (DEGs) obtained using Seurat’s *FindMarkers* command and known cell type-specific markers found in the literature.

### Genomic phylostratigraphy mapping of protein-coding genes in *Mus musculus* and *Danio rerio* genomes

The phylostratigraphic procedure was performed as previously described[40, 42, 45]. Consensus phylogenies covering divergence from the last common ancestor of cellular organisms to *M. musculus* and *D. rerio* as focal organisms were constructed following the recent phylogenetic literature[93–100]. The nodes were chosen based on their phylogenetic support from the literature, the availability of reference genomes for terminal taxa, and their importance in evolutionary transitions. The complete set of protein sequences for 513 (mouse) and 506 (zebrafish) terminal taxa was retrieved from Ensembl and NCBI databases. The protein sequence database was prepared for sequence similarity searches by checking the consistency of the files, leaving only the longest splicing variant per eukaryotic gene and adding taxon tags to sequence headers. The phylostratigraphy map of *M. musculus* and *D.rerio* was constructed by comparing 22,769 *M. musculus* and 25,787 *D.rerio* protein sequences with the protein sequence database by blastp algorithm V2.9.0 with a 10^-3^ e-value threshold[101]. Protein sequences that did not return their sequence as a match were discarded, which left 22,590 and 25,721 protein sequences in mouse and zebrafish, respectively. The mouse and zebrafish protein sequences were mapped on consensus phylogenies with 22 and 16 phylostrata (ps), respectively. Each protein was assigned to the oldest internode on the phylogeny, where it still had a match[40–42, 45].

### Transcriptome Age Index

We generated a normalized relative count matrix from the Seurat object using the total count normalization method multiplied by a scaling factor of 10^6^. The metadata was extracted from the Seurat file to provide the identities of each cell in the count matrix. The gene expression matrix was combined with the phylogenetic age assignment of genes from the phylostratigraphy map to calculate each cell’s TAI[45]. TAI was calculated using myTAI[102] R package based on the formula:

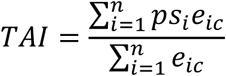

where the *ps_i_* represents the phylostratum assignment of gene *i* and *e_ic_* denotes the normalized read count of gene *i* in a cell (*c*), and *n* is the total number of expressed genes in a cell (*c*). The read count of each gene acts as a weight factor to give more substantial weight to highly expressed genes. We computed TAI values using all expressed genes and specifically expressed genes. To find the genes that are specifically upregulated in a cell type, we performed differential gene expression analysis using Seurat’s *FindAllMarkers* command. We only considered genes that show a minimum 25% difference in the fraction of detection (min.diff.pct= 0.25) between the focal cell type population and the rest of the cell populations. We then projected the TAI values of each cell according to their cell type identities to infer the relative age of each skeletal cell type. The analysis was repeated after the number of cells for each cell type was downsampled using Seurat’s *subset* function (n=50, n=100).

The TAI values of cell types were statistically compared using Kruskal-Wallis and pairwise Wilcoxon post hoc tests. *P* values were adjusted for multiple comparisons using Benjamini and Hochberg[103].

Partial TAI was calculated based on the alternative formula of TAI:

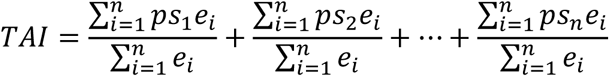

representing the sum of partial TAI concentrations split according to the phylogenetic rank, ps.

### Average and relative expression analysis

We calculated the average expression of genes for each ps and every individual cell using the normalized total counts multiplied by a scaling factor of 10^6^. The relative expression of genes for a given phylostratum and cell type was subsequently calculated by linearly transforming the average expression values of genes from each phylostratum and cell type using the formula[45]:

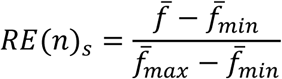

where *^f̅^* represents the average expression of genes from phylostratum n for cell type *s*, and *^f̅^_min_ and ^f̅^_max_* denote the minimum and maximum mean expression values from phylostratum n across all cell types, respectively.

### Enrichment analysis

The enrichment of individual cell types across phylostrata was performed using the two-tailed hypergeometric test by comparing the distribution of all differentially expressed genes (DEGs) for each cell type (test set) across phylostrata with the overall distribution of DEGs from cell types of mesenchymal lineage (background set). Deviations in distributions are presented with log-odds, where log-odds of zero denote that the observed frequency of DEGs in a phylostrata equals the expected frequency from the background set. Enrichments in certain phylostrata indicate that a significant number of genes are associated with the evolution of the cell type. The test was adjusted for multiple comparisons by correcting p-values using the Benjamini and Hochberg procedure[103].

### GO enrichment

Gene ontology (GO) enrichment analysis of gene sets was performed and visualized with clusterProfiler R package and the *enrichGO* function correcting for multiple comparisons using the Benjamini and Hochberg procedure[103]. Genome-wide *M. musculus* and *D. rerio* annotations were provided by the org.Mm.eg.db[104] and org.Dr.eg.db[105] R packages, respectively. All genes provided in the database were used as a background set.

## Ethical Statement

This study used previously published data from other laboratories. No new results were generated from live specimens in the course of this research.

## Data, Materials and Software Availability

Mouse single-cell dataset is available at NCBI (Geo accession number GSE142425). Zebrafish single-cell datasets are available at FaceBase (Record ID 5-DAQ4) and Zebrahub (https://zebrahub.ds.czbiohub.org). Phylostratigraphy data are presented in the supplementary information.

## Supplementary Material

This file contains the Supplementary information.

## Acknowledgments

AD is supported by the International Max Planck Research School for Evolutionary Biology (IMPRS-EB) of the Max Planck Society. MK is supported by the Max Planck Society. AK is supported by grants from the German Research Foundation (DFG) project KL 3475/2-1 and CRC 1461: “Neurotronics: Bio-Inspired Information Pathways”. The figures were created with BioRender.com.

## Author contributions

AD: Conceptualization, Methodology, Software, Formal analysis, Data Curation, Writing-Original Draft, Writing-Review and Editing, Visualization. SK: Methodology, Software, Formal Analysis, Writing-Review and Editing. KU: Methodology, Software, Writing-Review and Editing. TDL: Conceptualization, Methodology, Formal Analysis, Writing-Review and Editing. AK: Conceptualization, Formal Analysis, Writing-Review and Editing, Visualization, Supervision. MK: Conceptualization, Formal analysis, Writing-Original Draft, Writing-Review and Editing, Visualization, Supervision, Project administration.

## Competing interestss

The authors declare no conflict of interest.

