## Supplementary Information for "Evolutionary trends in the emergence of skeletal cell types"

### Supplementary Information
### Evolutionary trends in the emergence of skeletal cell types

Amor Damatac II, Sara Koska, Kristian K Ullrich, Tomislav Domazet-Lošo, Alexander Klimovich, Markéta Kaucká*

**This PDF file includes:**

Supplementary Result
Figures S1 to S9

**Other supporting materials for this manuscript include the following:**

Tables S1-18

#### Supplementary Result

##### Single-cell phylotranscriptomic analysis of skeletal cell types in zebrafish

In order to test whether the evolutionary trends of skeletal cell type emergence revealed in the mouse dataset can be recapitulated using single-cell data from another species, we searched for available datasets that would contain the spectrum of skeletal cell types. At the time of this investigation, we could take advantage of only two existing datasets. Here, we applied the same research strategy to skeletal cell types obtained from zebrafish (*Danio rerio*) developing larvae single-cell transcriptome analysis(1, 2). Using the same skeletal marker genes, we identified IC and OB in the Fabian dataset(1) at the stages five days post-fertilization (5dpf) and 14 days post-fertilization (14 dpf) (SI Appendix, Fig. S8B). IC is also present in Lange dataset(2) at 5dpf; however, we noticed that *col10a1a*-positive cells labeled by the authors as dermal bone clearly express a signature of both HC (*ihha*-positive) and OB (*col1a1a*-positive). Therefore, we assign these cells as a hybrid HC/OB cluster expressing both *ihha* and *col10a1* for the following analyses (Fig. S8B).

We performed genomic phylostratigraphy with zebrafish being the focal species to assign the phylogenetic age of 25,721 protein-coding genes across 16 ps (SI Appendix, Fig. S8A, S9; Tables S16-18).

Even though some cell types from the skeletogenic lineage are not present in the zebrafish dataset, the TAI profiling recapitulated patterns similar to those observed in mouse. OB and HC/OB have a higher TAI, which points at more recent evolutionary origin compared to IC (SI Appendix, Fig. S8C). Enrichment analyses show that zebrafish OB transcriptome is significantly associated with genes originated in Opisthokonta (ps3), and similarly to the OB in mouse in Holozoa (ps4) (SI Appendix, Fig. S8D). In this phylogenetic node, OB-related genes are significantly associated with biological processes involved in bone biology including ECM organization, skeletal development, protein kinase/ transferase activity and phosphorus/phosphate metabolism (SI Appendix, Fig. S8E). These include transcription factors from the Runx family, similar to mouse. For HC/OB, we detected enrichment signal in Holozoa (ps4), Metazoa (ps5), Bilateria (ps7), Vertebrata (ps11) and Gnathostosmata (ps12) (Fig. 3d), similar to HC and OB in mouse (SI Appendix, Fig. S8D). GO enrichment profiling of genes from the early phylostrata uncovered their association with diverse biological modules known to be involved in skeletogenesis (SI Appendix, Fig. S8F), similar to the GO terms we observed in mouse HC. Again, we recovered transcription factors from Runx family. Overall, these results in zebrafish corroborate our findings in mouse, and allow to infer the history of skeletal cell type emergence and evolution through recruitment of transcription factors and novel genes, enabling specific functions.

1. P. Fabian *et al.*, Lifelong single-cell profiling of cranial neural crest diversification in zebrafish. *Nat Commun* **13**, 13 (2022).
2.M. Lange *et al.*, Zebrahub – Multimodal Zebrafish Developmental Atlas Reveals the State Transition Dynamics of Late Vertebrate Pluripotent Axial Progenitors. *bioRxiv* https://doi.org/10.1101/2023.03.06.531398 (2023).

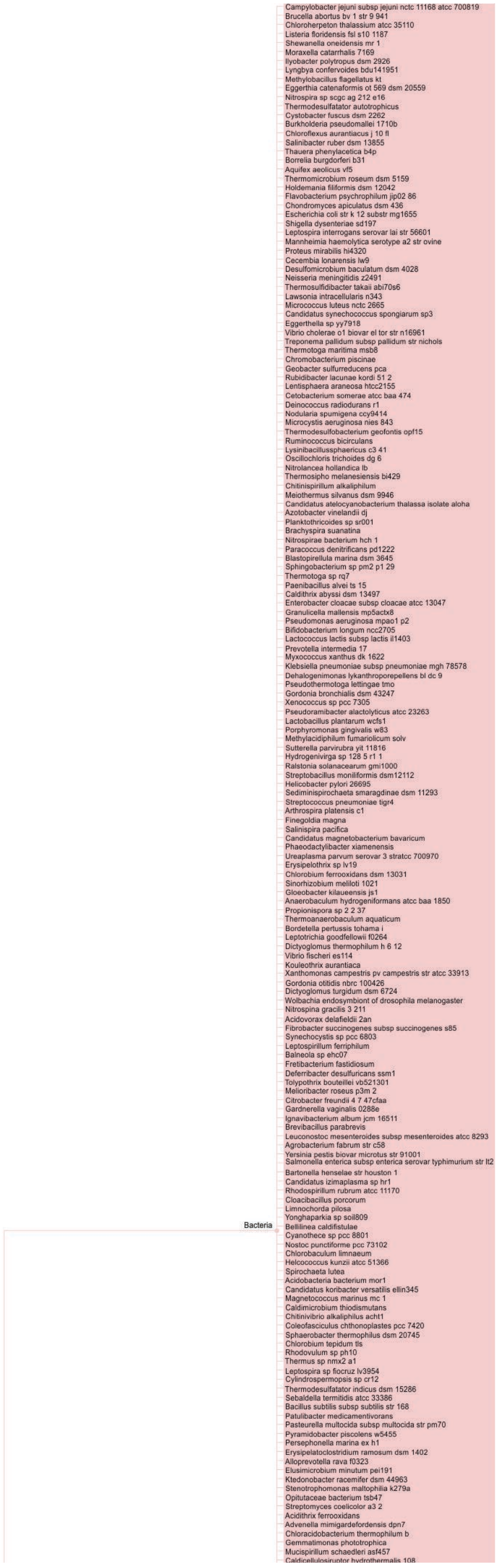

Campylobacter jejuni subsp jejuni nctc 11168 atcc 700819
Brucella abortus bv 1 str 9 941
Chloroherpeton thalassium atcc 35110
Listeria floridensis fsl s10 1187
Shewanella oneidensis mr 1
Moraxella catarrhalis 7169
Ilyobacter polytropus dsm 2926
Lyngbya confervoides bdu141951
Methylobacillus flagellatus kt
Eggerthia catenaformis ot 569 dsm 20559
Nitrospira sp scgc ag 212 e16
Thermodesulfatator autotrophicus
Cystobacter fuscus dsm 2262
Burkholderia pseudomallei 1710b
Chloroflexus aurantiacus j 10 fl
Salinibacter ruber dsm 13855
Thauera phenylacetica b4p
Borrelia burgdorferi b31
Aquifex aeolicus vf5
Thermomicrobium roseum dsm 5159
Holdemania filiformis dsm 12042
Flavobacterium psychrophilum jip02 86
Chondromyces apiculatus dsm 436
Escherichia coli str k 12 substr mg1655
Shigella dysenteriae sd197
Leptospira interrogans serovar lai str 56601
Mannheimia haemolytica serotype a2 str ovine
Proteus mirabilis hi4320
Cecembia lonarensis lw9
Desulfomicrobium baculatum dsm 4028
Neisseria meningitidis z2491
Thermosulfidibacter takaii abi70s6
Lawsonia intracellularis n343
Micrococcus luteus nctc 2665
Candidatus synechococcus spongiarum sp3
Eggerthella sp yy7918
Vibrio cholerae o1 biovar el tor str n16961
Treponema pallidum subsp pallidum str nichols
Thermotoga maritima msb8
Chromobacterium piscinae
Geobacter sulfurreducens pca
Rubidibacter lacunae kordi 51 2
Lentisphaera araneosa htcc2155
Cetobacterium somerae atcc baa 474
Deinococcus radiodurans r1
Nodularia spumigena ccy9414
Microcystis aeruginosa nies 843
Thermodesulfobacterium geofontis opf15
Ruminococcus bicirculans
Lysinibacillussphaericus c3 41
Oscillochloris trichoides dg 6
Nitrolancea hollandica lb
Thermosipho melanesiensis bi429
Chitinispirillum alkaliphilum
Meiothermus silvanus dsm 9946
Candidatus atelocyanobacterium thalassa isolate aloha
Azotobacter vinelandii dj
Planktothricoides sp sr001
Brachyspira suanatina
Nitrospirae bacterium hch 1
Paracoccus denitrificans pd1222
Blastopirellula marina dsm 3645
Sphingobacterium sp pm2 p1 29
Thermotoga sp rq7
Paenibacillus alvei ts 15
Caldithrix abyssi dsm 13497
Enterobacter cloacae subsp cloacae atcc 13047
Granulicella mallensis mp5actx8
Pseudomonas aeruginosa mpao1 p2
Bifidobacterium longum ncc2705
Lactococcus lactis subsp lactis il1403
Prevotella intermedia 17
Myxococcus xanthus dk 1622
Klebsiella pneumoniae subsp pneumoniae mgh 78578
Dehalogenimonas lykanthroporepellens bl dc 9
Pseudothermotoga lettingae tmo
Gordonia bronchialis dsm 43247
Xenococcus sp pcc 7305
Pseudoramibacter alactolyticus atcc 23263
Lactobacillus plantarum wcfs1
Porphyromonas gingivalis w83
Methylacidiphilum fumariolicum solv
Sutterella parvirubra yit 11816
Hydrogenivirga sp 128 5 r1 1
Ralstonia solanacearum gmi1000
Streptobacillus moniliformis dsm12112
Helicobacter pylori 26695
Sediminispirochaeta smaragdinae dsm 11293
Streptococcus pneumoniae tigr4
Arthrospira platensis c1
Finegoldia magna
Salinispira pacifica
Candidatus magnetobacterium bavaricum
Phaeodactylibacter xiamenensis
Ureaplasma parvum serovar 3 stratcc 700970
Erysipelothrix sp lv19
Chlorobium ferrooxidans dsm 13031
Sinorhizobium meliloti 1021
Gloeobacter kilaueensis js1
Anaerobaculum hydrogeniformans atcc baa 1850
Propionispora sp 2 2 37
Thermoanaerobaculum aquaticum
Bordetella pertussis tohama i
Leptotrichia goodfellowii f0264
Dictyoglomus thermophilum h 6 12
Vibrio fischeri es114
Kouleothrix aurantiaca
Xanthomonas campestris pv campestris str atcc 33913
Gordonia otitidis nbrc 100426
Dictyoglomus turgidum dsm 6724
Wolbachia endosymbiont of drosophila melanogaster
Nitrospina gracilis 3 211
Acidovorax delafieldii 2an
Fibrobacter succinogenes subsp succinogenes s85
Synechocystis sp pcc 6803
Leptospirillum ferriphilum
Balneola sp ehc07
Fretibacterium fastidiosum
Deferribacter desulfuricans ssm1
Tolypothrix bouteillei vb521301
Melioribacter roseus p3m 2
Citrobacter freundii 4 7 47cfaa
Gardnerella vaginalis 0288e
Ignavibacterium album jcm 16511
Brevibacillus parabrevis
Leuconostoc mesenteroides subsp mesenteroides atcc 8293
Agrobacterium fabrum str c58
Yersinia pestis biovar microtus str 91001
Salmonella enterica subsp enterica serovar typhimurium str lt2
Bartonella henselae str houston 1
Candidatus izimaplasma sp hr1
Rhodospirillum rubrum atcc 11170
Cloacibacillus porcorum
Limnochorda pilosa
Yonghaparkia sp soil809
Bacteria
Bellilinea caldifistulae
Cyanothece sp pcc 8801
Nostoc punctiforme pcc 73102
Chlorobaculum limnaeum
Helcococcus kunzii atcc 51366
Spirochaeta lutea
Acidobacteria bacterium mor1
Candidatus koribacter versatilis ellin345
Magnetococcus marinus mc 1
Caldimicrobium thiodismutans
Chitinivibrio alkaliphilus acht1
Coleofasciculus chthonoplastes pcc 7420
Sphaerobacter thermophilus dsm 20745
Chlorobium tepidum tls
Rhodovulum sp ph10
Thermus sp nmx2 a1
Leptospira sp fiocruz lv3954
Cylindrospermopsis sp cr12
Thermodesulfatator indicus dsm 15286
Sebaldella termitidis atcc 33386
Bacillus subtilis subsp subtilis str 168
Patulibacter medicamentivorans
Pasteurella multocida subsp multocida str pm70
Pyramidobacter piscolens w5455
Persephonella marina ex h1
Erysipelatoclostridium ramosum dsm 1402
Alloprevotella rava f0323
Elusimicrobium minutum pei191
Ktedonobacter racemifer dsm 44963
Stenotrophomonas maltophilia k279a
Opitutaceae bacterium tsb47
Streptomyces coelicolor a3 2
Acidithrix ferrooxidans
Advenella mimigardefordensis dpn7
Chloracidobacterium thermophilum b
Gemmatimonas phototrophica
Mucispirillum schaedleri asf457
Caldicellulosiruptor hydrothermalis 108

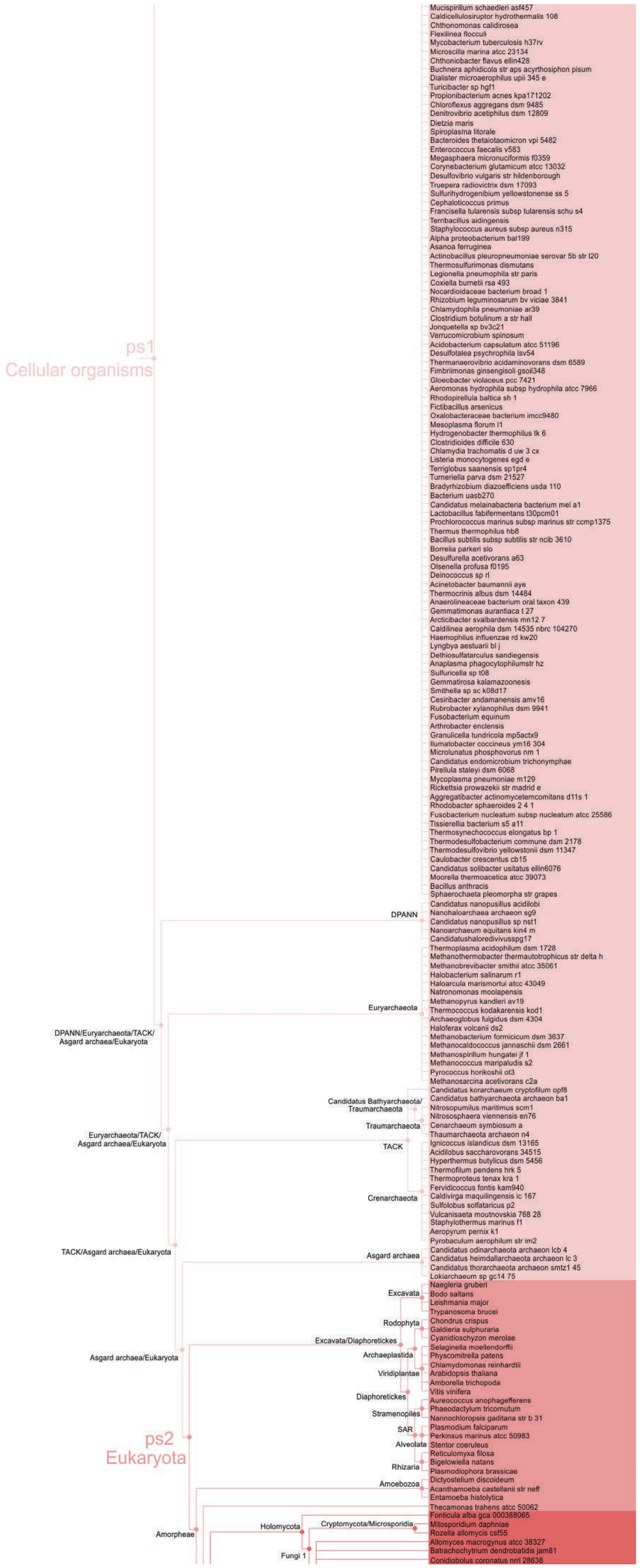

ps1
Cellular organisms
Mucispirillum schaedleri asf457
Caldicellulosiruptor hydrothermalis 108
Chthonomonas calidirosea
Flexilinea flocculi
Mycobacterium tuberculosis h37rv
Microscilla marina atcc 23134
Chthoniobacter flavus ellin428
Buchnera aphidicola str aps acyrthosiphon pisum
Dialister microaerophilus upii 345 e
Turicibacter sp hgf1
Propionibacterium acnes kpa171202
Chloroflexus aggregans dsm 9485
Denitrovibrio acetiphilus dsm 12809
Dietzia maris
Spiroplasma litorale
Bacteroides thetaiotaomicron vpi 5482
Enterococcus faecalis v583
Megasphaera micronuciformis f0359
Corynebacterium glutamicum atcc 13032
Desulfovibrio vulgaris str hildenborough
Truepera radiovictrix dsm 17093
Sulfurihydrogenibium yellowstonense ss 5
Cephaloticoccus primus
Francisella tularensis subsp tularensis schu s4
Terribacillus aidingensis
Staphylococcus aureus subsp aureus n315
Alpha proteobacterium bal199
Asanoa ferruginea
Actinobacillus pleuropneumoniae serovar 5b str l20
Thermosulfurimonas dismutans
Legionella pneumophila str paris
Coxiella burnetii rsa 493
Nocardioidaceae bacterium broad 1
Rhizobium leguminosarum bv viciae 3841
Chlamydophila pneumoniae ar39
Clostridium botulinum a str hall
Jonquetella sp bv3c21
Verrucomicrobium spinosum
Acidobacterium capsulatum atcc 51196
Desulfotalea psychrophila lsv54
Thermanaerovibrio acidaminovorans dsm 6589
Fimbriimonas ginsengisoli gsoil348
Gloeobacter violaceus pcc 7421
Aeromonas hydrophila subsp hydrophila atcc 7966
Rhodopirellula baltica sh 1
Fictibacillus arsenicus
Oxalobacteraceae bacterium imcc9480
Mesoplasma florum l1
Hydrogenobacter thermophilus tk 6
Clostridioides difficile 630
Chlamydia trachomatis d uw 3 cx
Listeria monocytogenes egd e
Terriglobus saanensis sp1pr4
Turneriella parva dsm 21527
Bradyrhizobium diazoefficiens usda 110
Bacterium uasb270
Candidatus melainabacteria bacterium mel a1
Lactobacillus fabifermentans t30pcm01
Prochlorococcus marinus subsp marinus str ccmp1375
Thermus thermophilus hb8
Bacillus subtilis subsp subtilis str ncib 3610
Borrelia parkeri slo
Desulfurella acetivorans a63
Olsenella profusa f0195
Deinococcus sp rl
Acinetobacter baumannii aye
Thermocrinis albus dsm 14484
Anaerolineaceae bacterium oral taxon 439
Gemmatimonas aurantiaca t 27
Arcticibacter svalbardensis mn12 7
Caldilinea aerophila dsm 14535 nbrc 104270
Haemophilus influenzae rd kw20
Lyngbya aestuarii bl j
Dethiosulfatarculus sandiegensis
Anaplasma phagocytophilumstr hz
Sulfuricella sp t08
Gemmatirosa kalamazoonesis
Smithella sp sc k08d17
Cesiribacter andamanensis amv16
Rubrobacter xylanophilus dsm 9941
Fusobacterium equinum
Arthrobacter enclensis
Granulicella tundricola mp5actx9
Ilumatobacter coccineus ym16 304
Microlunatus phosphovorus nm 1
Candidatus endomicrobium trichonymphae
Pirellula staleyi dsm 6068
Mycoplasma pneumoniae m129
Rickettsia prowazekii str madrid e
Aggregatibacter actinomycetemcomitans d11s 1
Rhodobacter sphaeroides 2 4 1
Fusobacterium nucleatum subsp nucleatum atcc 25586
Tissierella bacterium s5 a11
Thermosynechococcus elongatus bp 1
Thermodesulfobacterium commune dsm 2178
Thermodesulfovibrio yellowstonii dsm 11347
Caulobacter crescentus cb15
Candidatus solibacter usitatus ellin6076
Moorella thermoacetica atcc 39073
Bacillus anthracis
Sphaerochaeta pleomorpha str grapes
DPANN
Candidatus nanopusillus acidilobi
Nanohaloarchaea archaeon sg9
Candidatus nanopusillus sp nst1
Nanoarchaeum equitans kin4 m
Candidatushaloredivivussg17
Thermoplasma acidophilum dsm 1728
Methanothermobacter thermautotrophicus str delta h
Methanobrevibacter smithii atcc 35061
Halobacterium salinarum r1
Haloarcula marismortui atcc 43049
Natronomonas moolapensis
Methanopyrus kandleri av19
Euryarchaeota
Thermococcus kodakarensis kod1
Archaeoglobus fulgidus dsm 4304
DPANN/Euryarchaeota/TACK/
Asgard archaea/Eukaryota
Haloferax volcanii ds2
Methanobacterium formicicum dsm 3637
Methanocaldococcus jannaschii dsm 2661
Methanospirillum hungatei jf 1
Methanococcus maripaludis s2
Pyrococcus horikoshii ot3
Methanosarcina acetivorans c2a
Candidatus korarchaeum cryptofilum opf8
Candidatus Bathyarchaeota/
Traumarchaeota
Candidatus bathyarchaeota archaeon ba1
Nitrosopumilus maritimus scm1
Nitrososphaera viennensis en76
Traumarchaeota
Cenarchaeum symbiosum a
Euryarchaeota/TACK/
Asgard archaea/Eukaryota
Thaumarchaeota archaeon n4
TACK
Ignicoccus islandicus dsm 13165
Acidilobus saccharovorans 34515
Hyperthermus butylicus dsm 5456
Thermofilum pendens hrk 5
Thermoproteus tenax kra 1
Fervidicoccus fontis kam940
Crenarchaeota
Caldivirga maquilingensis ic 167
Sulfolobus solfataricus p2
Vulcanisaeta moutnovskia 768 28
Staphylothermus marinus f1
Aeropyrum pernix k1
Pyrobaculum aerophilum str im2
TACK/Asgard archaea/Eukaryota
Candidatus odinarchaeota archaeon lcb 4
Asgard archaea
Candidatus heimdallarchaeota archaeon lc 3
Candidatus thorarchaeota archaeon smtz1 45
Lokiarchaeum sp gc14 75
Naegleria gruberi
Excavata
Bodo saltans
Leishmania major
Trypanosoma brucei
Rodophyta
Chondrus crispus
Galdieria sulphuraria
Excavata/Diaphoretickes
Cyanidioschyzon merolae
Asgard archaea/Eukaryota
Selaginella moellendorffii
Archaeplastida
Physcomitrella patens
Chlamydomonas reinhardtii
Viridiplantae
Arabidopsis thaliana
Amborella trichopoda
Vitis vinifera
Diaphoretickes
Aureococcus anophagefferens
Phaeodactylum tricornutum
Stramenopiles
Nannochloropsis gaditana str b 31
SAR
Plasmodium falciparum
ps2
Eukaryota
Perkinsus marinus atcc 50983
Alveolata
Stentor coeruleus
Reticulomyxa filosa
Rhizaria
Bigelowiella natans
Plasmodiophora brassicae
Dictyostelium discoideum
Amoebozoa
Acanthamoeba castellanii str neff
Entamoeba histolytica
Thecamonas trahens atcc 50062
Fonticula alba gca 000388065
Holomycota
Cryptomycota/Microsporidia
Mitosporidium daphniae
Amorpheae
Rozella allomycis csf55
Allomyces macrogynus atcc 38327
Batrachochytrium dendrobatidis jam81
Fungi 1
Conidiobolus coronatus nrrl 28638

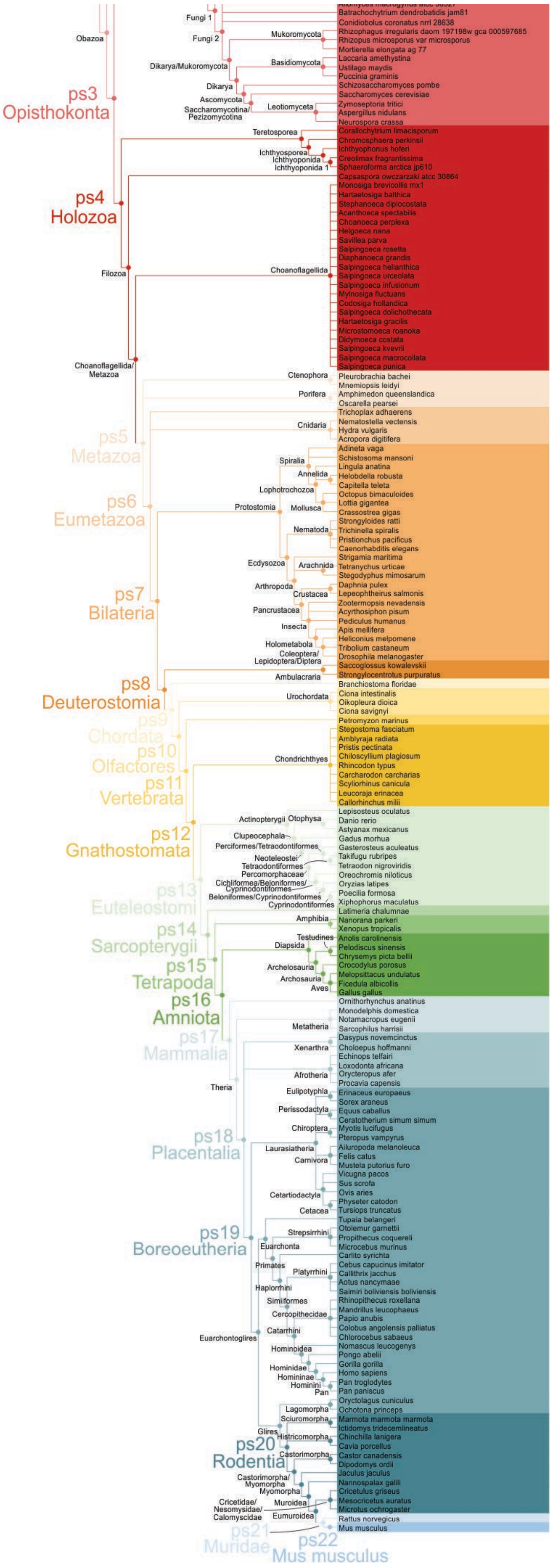

Batrachochytrium dendrobatidis jam81
Conidiobolus coronatus nrrl 28638
Rhizophagus irregularis daom 197198w gca 000597685
Rhizopus microsporus var microsporus
Mortierella elongata ag 77
Laccaria amethystina
Ustilago maydis
Puccinia graminis
Schizosaccharomyces pombe
Saccharomyces cerevisiae
Zymoseptoria tritici
Aspergillus nidulans
Neurospora crassa
Fungi 1
Fungi 2
Obazoa
Mukoromycota
Basidiomycota
Dikarya/Mukoromycota
Dikarya
Ascomycota
Saccharomycotina/
Pezizomycotina
Leotiomyceta
ps3
Opisthokonta
Corallochytrium limacisporum
Chromosphaera perkinsii
Ichthyophonus hoferi
Creolimax fragrantissima
Sphaeroforma arctica jp610
Capsaspora owczarzaki atcc 30864
Teretosporea
Ichthyosporea
Ichthyoponida
Ichthyoponida 1
ps4
Holozoa
Monosiga brevicollis mx1
Hartaetosiga balthica
Stephanoeca diplocostata
Acanthoeca spectabilis
Choanoeca perplexa
Helgoeca nana
Savillea parva
Salpingoeca rosetta
Diaphanoeca grandis
Salpingoeca helianthica
Salpingoeca urceolata
Salpingoeca infusionum
Mylnosiga fluctuans
Codosiga hollandica
Salpingoeca dolichothecata
Hartaetosiga gracilis
Microstomoeca roanoka
Didymoeca costata
Salpingoeca kvevrii
Salpingoeca macrocollata
Salpingoeca punica
Filozoa
Choanoflagellida
Choanoflagellida/
Metazoa
Ctenophora
Pleurobrachia bachei
Mnemiopsis leidyi
Porifera
Amphimedon queenslandica
Oscarella pearsei
Trichoplax adhaerens
Nematostella vectensis
Cnidaria
Hydra vulgaris
Acropora digitifera
ps5
Metazoa
Adineta vaga
Schistosoma mansoni
Lingula anatina
Helobdella robusta
Capitella teleta
Octopus bimaculoides
Lottia gigantea
Crassostrea gigas
Strongyloides ratti
Trichinella spiralis
Pristionchus pacificus
Caenorhabditis elegans
Strigamia maritima
Tetranychus urticae
Stegodyphus mimosarum
Daphnia pulex
Lepeophtheirus salmonis
Zootermopsis nevadensis
Acyrthosiphon pisum
Pediculus humanus
Apis mellifera
Heliconius melpomene
Tribolium castaneum
Drosophila melanogaster
Spiralia
Annelida
Lophotrochozoa
Protostomia
Mollusca
Nematoda
Ecdysozoa
Arachnida
Arthropoda
Crustacea
Pancrustacea
Insecta
Holometabola
Coleoptera/
Lepidoptera/Diptera
ps6
Eumetazoa
ps7
Bilateria
Saccoglossus kowalevskii
Strongylocentrotus purpuratus
Ambulacraria
ps8
Deuterostomia
Branchiostoma floridae
Ciona intestinalis
Oikopleura dioica
Ciona savignyi
Urochordata
Petromyzon marinus
ps9
Chordata
ps10
Olfactores
ps11
Vertebrata
Stegostoma fasciatum
Amblyraja radiata
Pristis pectinata
Chiloscyllium plagiosum
Rhincodon typus
Carcharodon carcharias
Scyliorhinus canicula
Leucoraja erinacea
Callorhinchus milii
Chondrichthyes
ps12
Gnathostomata
Lepisosteus oculatus
Danio rerio
Astyanax mexicanus
Gadus morhua
Gasterosteus aculeatus
Takifugu rubripes
Tetraodon nigroviridis
Oreochromis niloticus
Oryzias latipes
Poecilia formosa
Xiphophorus maculatus
Actinopterygii
Otophysa
Clupeocephala
Perciformes/Tetraodontiformes
Neoteleostei
Tetraodontiformes
Percomorphaceae
Cichliformea/Beloniformes/
Cyprinodontiformes
Beloniformes/Cyprinodontiformes
Cyprinodontiformes
ps13
Euteleostomi
Latimeria chalumnae
Amphibia
Nanorana parkeri
Xenopus tropicalis
ps14
Sarcopterygii
Anolis carolinensis
Pelodiscus sinensis
Chrysemys picta bellii
Crocodylus porosus
Melopsittacus undulatus
Ficedula albicollis
Gallus gallus
Testudines
Diapsida
Archelosauria
Archosauria
Aves
ps15
Tetrapoda
ps16
Amniota
Ornithorhynchus anatinus
Monodelphis domestica
Notamacropus eugenii
Sarcophilus harrisii
Metatheria
ps17
Mammalia
Dasypus novemcinctus
Choloepus hoffmanni
Echinops telfairi
Loxodonta africana
Orycteropus afer
Procavia capensis
Xenarthra
Afrotheria
Theria
Erinaceus europaeus
Sorex araneus
Equus caballus
Ceratotherium simum simum
Myotis lucifugus
Pteropus vampyrus
Ailuropoda melanoleuca
Felis catus
Mustela putorius furo
Vicugna pacos
Sus scrofa
Ovis aries
Physeter catodon
Tursiops truncatus
Eulipotyphla
Perissodactyla
Chiroptera
Laurasiatheria
Carnivora
Cetartiodactyla
Cetacea
ps18
Placentalia
Tupaia belangeri
Otolemur garnettii
Propithecus coquereli
Microcebus murinus
Carlito syrichta
Cebus capucinus imitator
Callithrix jacchus
Aotus nancymaae
Saimiri boliviensis boliviensis
Rhinopithecus roxellana
Mandrillus leucophaeus
Papio anubis
Colobus angolensis palliatus
Chlorocebus sabaeus
Nomascus leucogenys
Pongo abelii
Gorilla gorilla
Homo sapiens
Pan troglodytes
Pan paniscus
Strepsirrhini
Euarchonta
Primates
Platyrrhini
Haplorrhini
Simiiformes
Cercopithecidae
Catarrhini
Euarchontoglires
Hominoidea
Hominidae
Homininae
Hominini
Pan
ps19
Boreoeutheria
Oryctolagus cuniculus
Ochotona princeps
Marmota marmota marmota
Ictidomys tridecemlineatus
Chinchilla lanigera
Cavia porcellus
Castor canadensis
Dipodomys ordii
Jaculus jaculus
Nannospalax galili
Cricetulus griseus
Mesocricetus auratus
Microtus ochrogaster
Lagomorpha
Sciuromorpha
Glires
Histricomorpha
Castorimorpha
ps20
Rodentia
Castorimorpha/
Myomorpha
Myomorpha
Cricetidae/
Nesomysidae/
Calomyscidae
Muroidea
Eumuroidea
Rattus norvegicus
Mus musculus
ps21
Muridae
ps22
Mus musculus

**Figure S1. Expanded consensus phylogeny used in the genomic phylostratigraphy analysis.** The tree covers divergence from the last common ancestor to mouse, *Mus musculus.* Twenty two nodes (phylostrata, ps) were considered in the analysis.

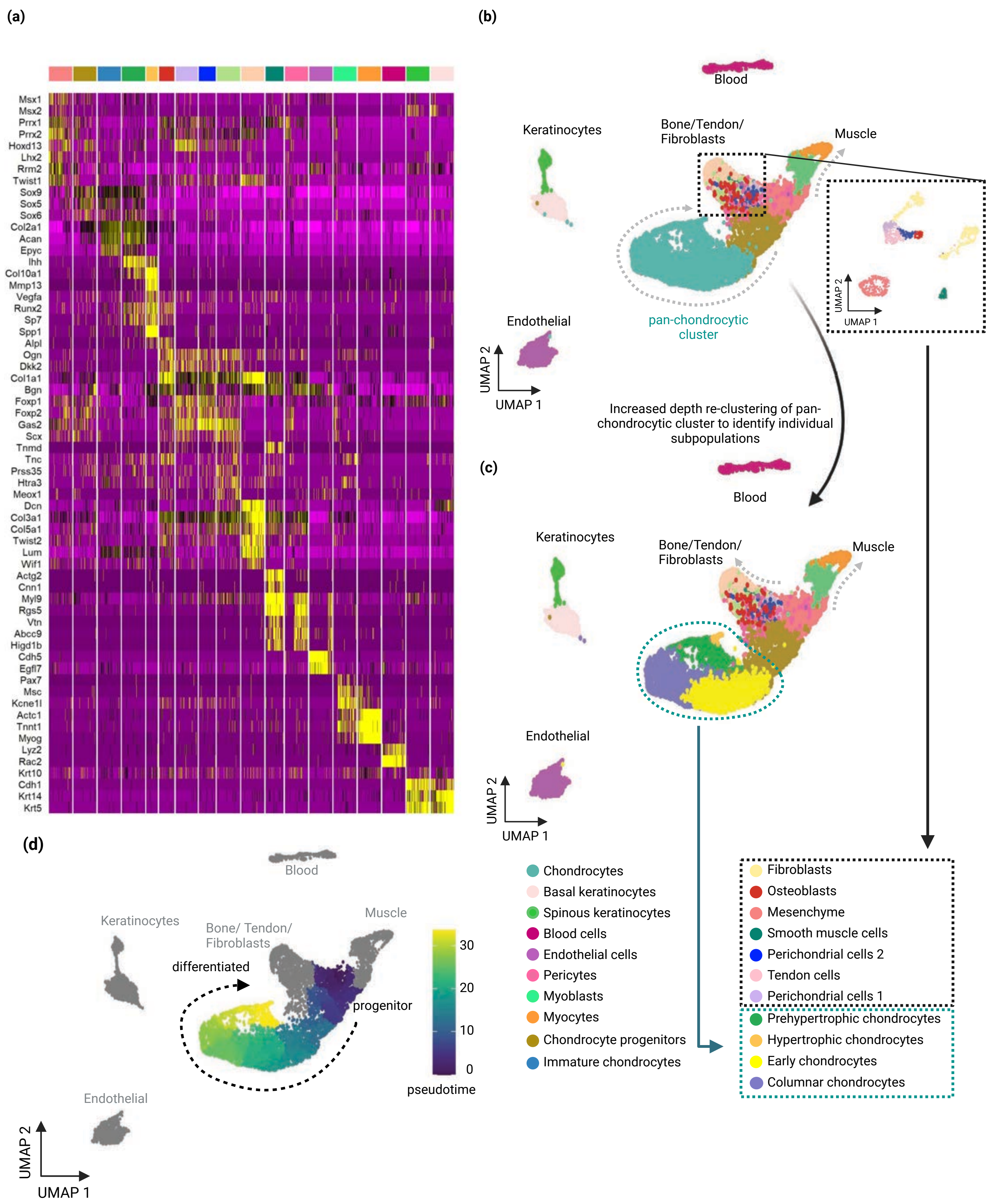


**Figure S2. Single cell transcriptome analysis.** (a) Gene expression profile of all cell types in the dataset using cell-type specific markers. (b,c)Two-dimensional UMAP of cell clusters shows the different cartilage populations at various clustering resolutions and subclustering of bone/tendon/fibroblast populations. (d) Monocle pseudotime trajectory of chondrogenic lineage cells starting undifferentiated mesenchyme to terminally differentiated HCs.

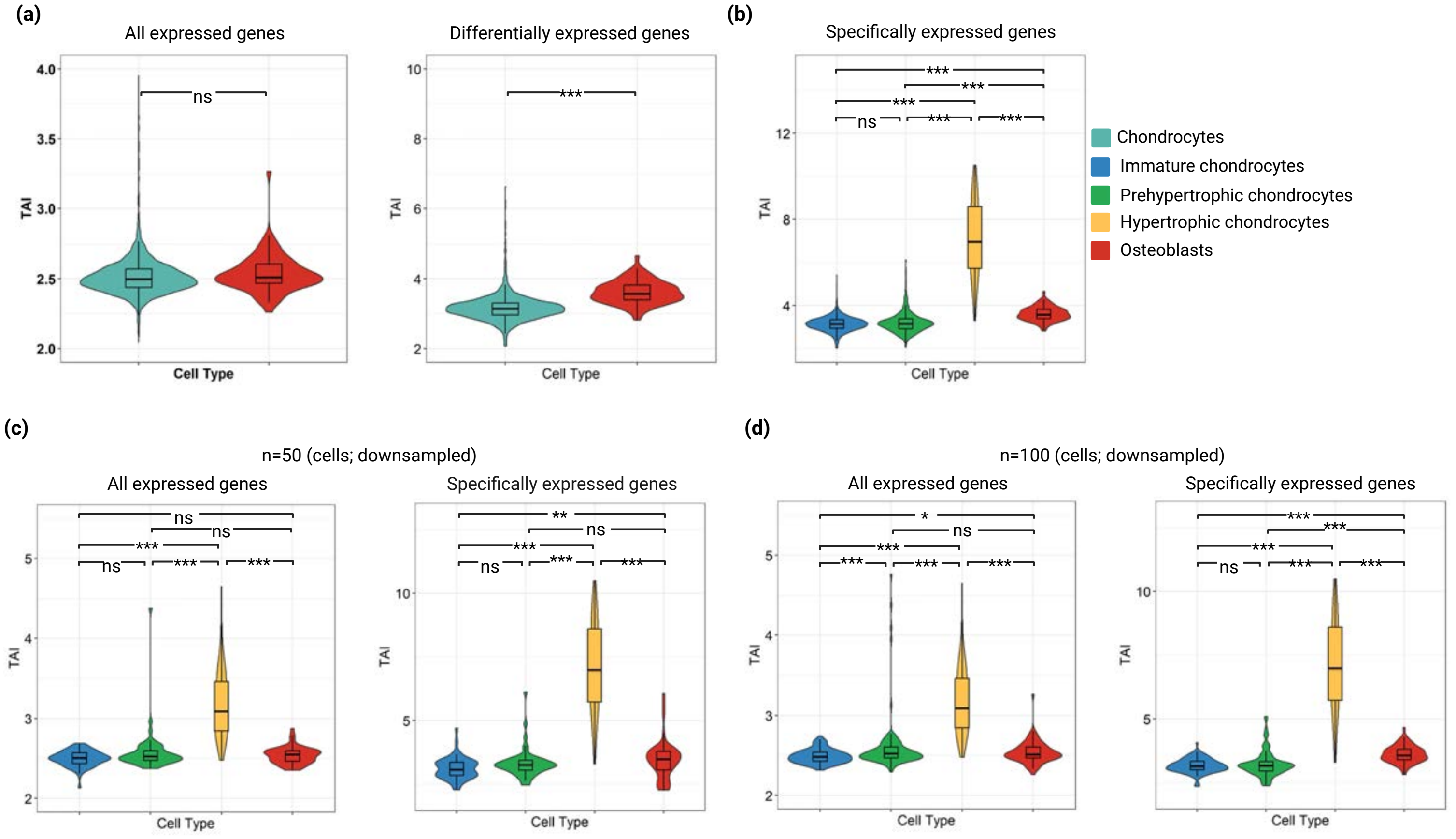


**Figure S3. Phylogenetic age of the cell types based on the transcriptome age index (TAI).** (a) TAI profile of chondrogenic and osteogenic cells. (b) TAI profile of IC, PHC, HC, and OC using specifically expressed genes (c) TAI profile of IC, PHC, HC, and OC after downsampling (n=50). (d) TAI profile of IC, PHC, HC, and OC after downsampling (n=100).

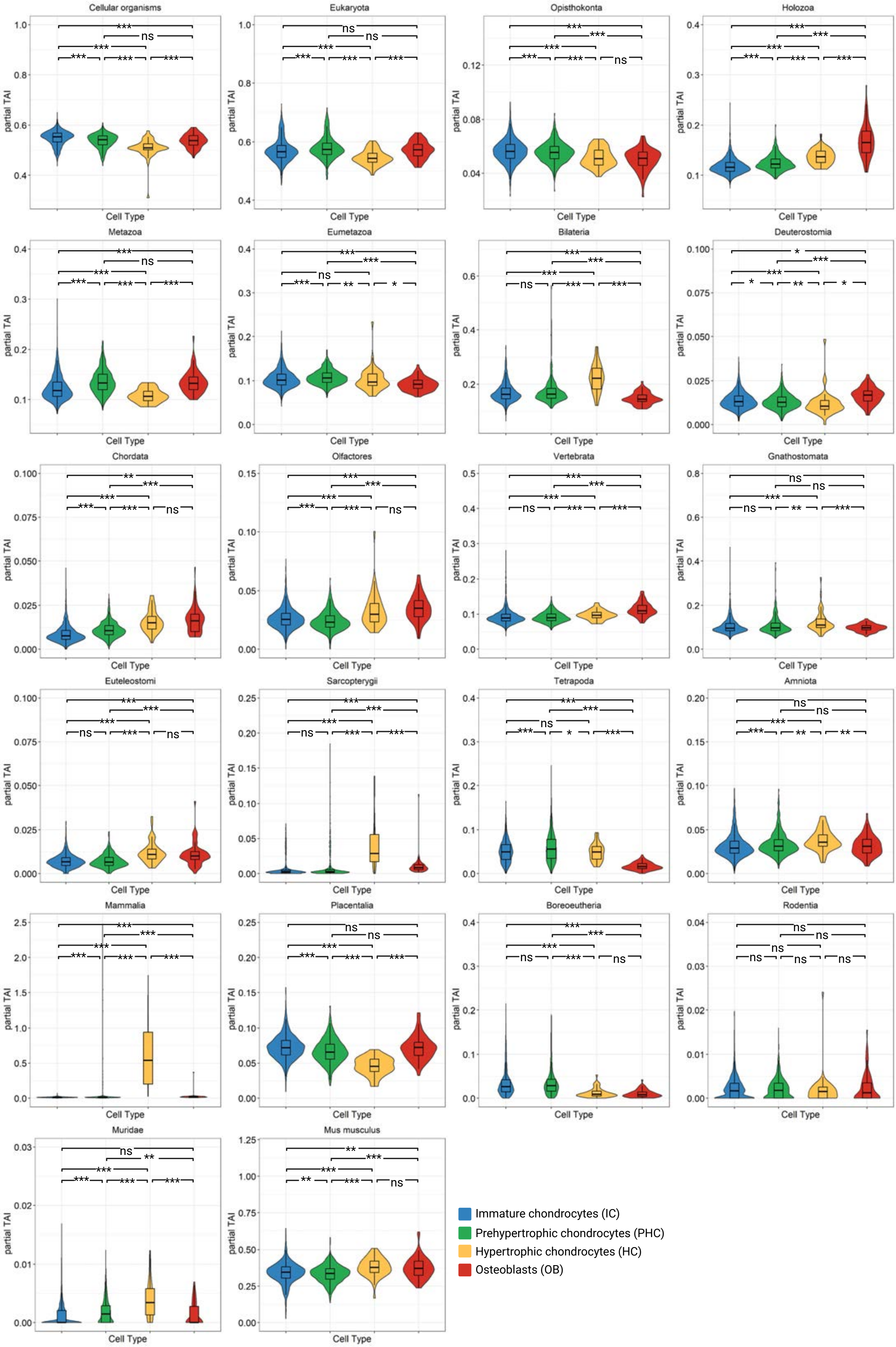
Cellular organisms
Eukaryota
Opisthokonta
Holozoa
Metazoa
Eumetazoa
Bilateria
Deuterostomia
Chordata
Olfactores
Vertebrata
Gnathostomata
Euteleostomi
Sarcopterygii
Tetrapoda
Amniota
Mammalia
Placentalia
Boreoeutheria
Rodentia
Muridae
Mus musculus
partial TAI
Cell Type
Immature chondrocytes (IC)
Prehypertrophic chondrocytes (PHC)
Hypertrophic chondrocytes (HC)
Osteoblasts (OB)

**Figure S4. Partial TAI (pTAI) of skeletal cell types according to the origin of genes from the different phylostrata**. Plots indicate the contribution of the different phylostrata to the global TAI. Statistical significance of differences among TAI values of skeletal cell types was evaluated using a pairwise Wilcoxon test corrected for multiple comparisons by BH. Asterisks denote adjusted p value levels (* ≤0.05, **≤0.01, ***≤0.001).

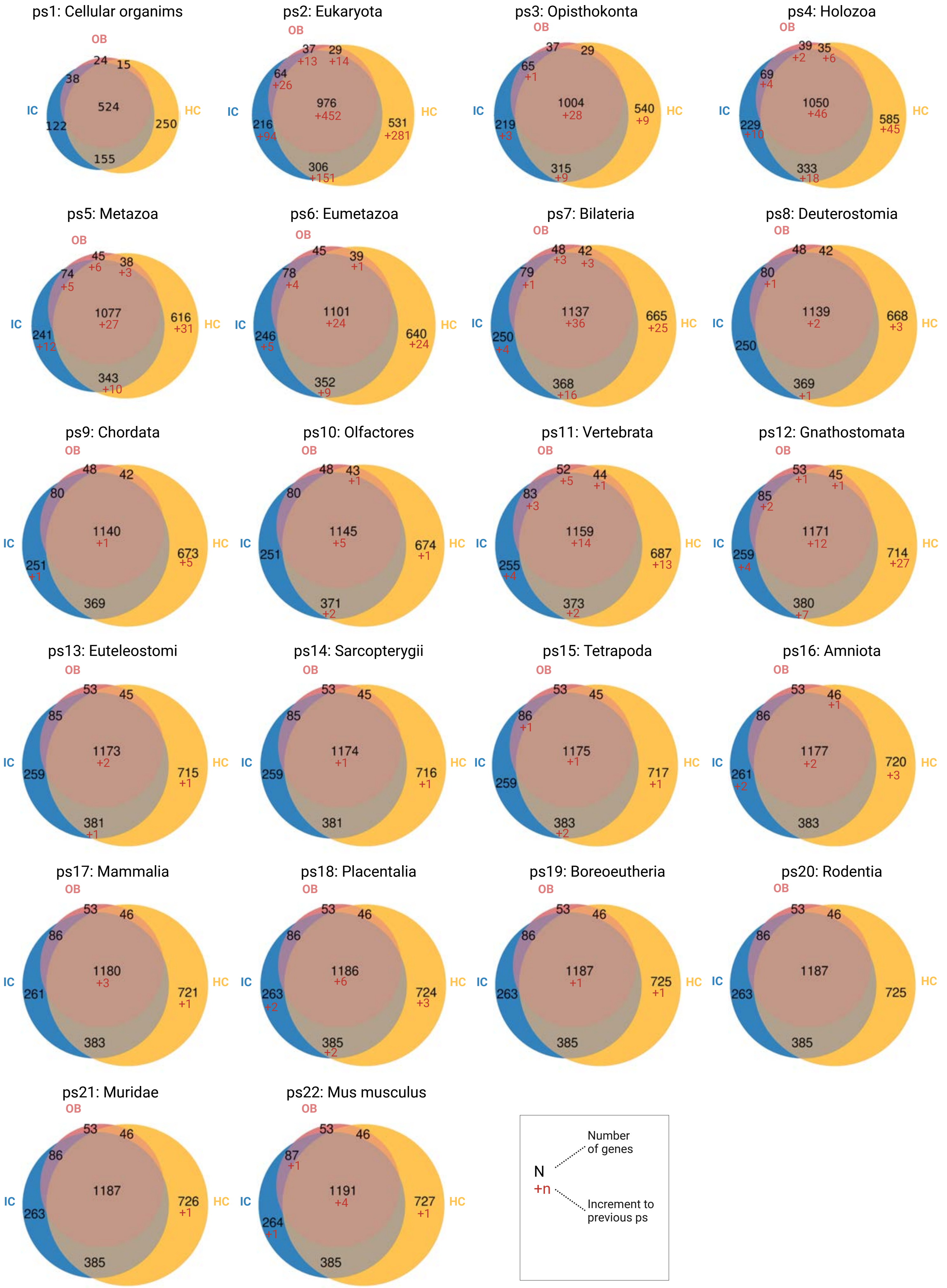
ps1: Cellular organims
OB IC HC
24 15 38 524 122 250 155
ps2: Eukaryota
OB IC HC
37 +13 29 +14 64 +26 976 +452 216 +94 531 +281 306 +151
ps3: Opisthokonta
OB IC HC
37 29 65 +1 1004 +28 540 +9 219 +3 315 +9
ps4: Holozoa
OB IC HC
39 +2 35 +6 69 +4 1050 +46 229 +10 585 +45 333 +18
ps5: Metazoa
OB IC HC
45 +6 38 +3 74 +5 1077 +27 616 +31 241 +12 343 +10
ps6: Eumetazoa
OB IC HC
45 39 +1 78 +4 1101 +24 246 +5 640 +24 352 +9
ps7: Bilateria
OB IC HC
48 +3 42 +3 79 +1 1137 +36 665 +25 250 +4 368 +16
ps8: Deuterostomia
OB IC HC
48 42 80 +1 1139 +2 668 +3 250 369 +1
ps9: Chordata
OB IC HC
48 42 80 1140 +1 251 +1 673 +5 369
ps10: Olfactores
OB IC HC
48 43 +1 80 1145 +5 251 674 +1 371 +2
ps11: Vertebrata
OB IC HC
52 +5 44 +1 83 +3 1159 +14 255 +4 687 +13 373 +2
ps12: Gnathostomata
OB IC HC
53 +1 45 +1 85 +2 1171 +12 259 +4 714 +27 380 +7
ps13: Euteleostomi
OB IC HC
53 45 85 1173 +2 259 715 +1 381 +1
ps14: Sarcopterygii
OB IC HC
53 45 85 1174 +1 259 716 +1 381
ps15: Tetrapoda
OB IC HC
53 45 86 +1 1175 +1 259 717 +1 383 +2
ps16: Amniota
OB IC HC
53 46 +1 86 1177 +2 261 +2 720 +3 383
ps17: Mammalia
OB IC HC
53 46 86 1180 +3 261 721 +1 383
ps18: Placentalia
OB IC HC
53 46 86 1186 +6 263 +2 724 +3 385 +2
ps19: Boreoeutheria
OB IC HC
53 46 86 1187 +1 263 725 +1 385
ps20: Rodentia
OB IC HC
53 46 86 1187 263 725 385
ps21: Muridae
OB IC HC
53 46 86 1187 263 726 +1 385
ps22: Mus musculus
OB IC HC
53 46 87 +1 1191 +4 264 +1 727 +1 385
N
+n
Number of genes
Increment to previous ps

**Figure S5. The proportions of shared and unique molecular components on the IC, HC and OC transcriptomes along the phylogeny.** Venn diagrams show the extent of shared and unique gene in the transcriptome of skeletal cell types. Only genes that were expressed by at least 50% of the cells in the given cell type were included in the analysis.

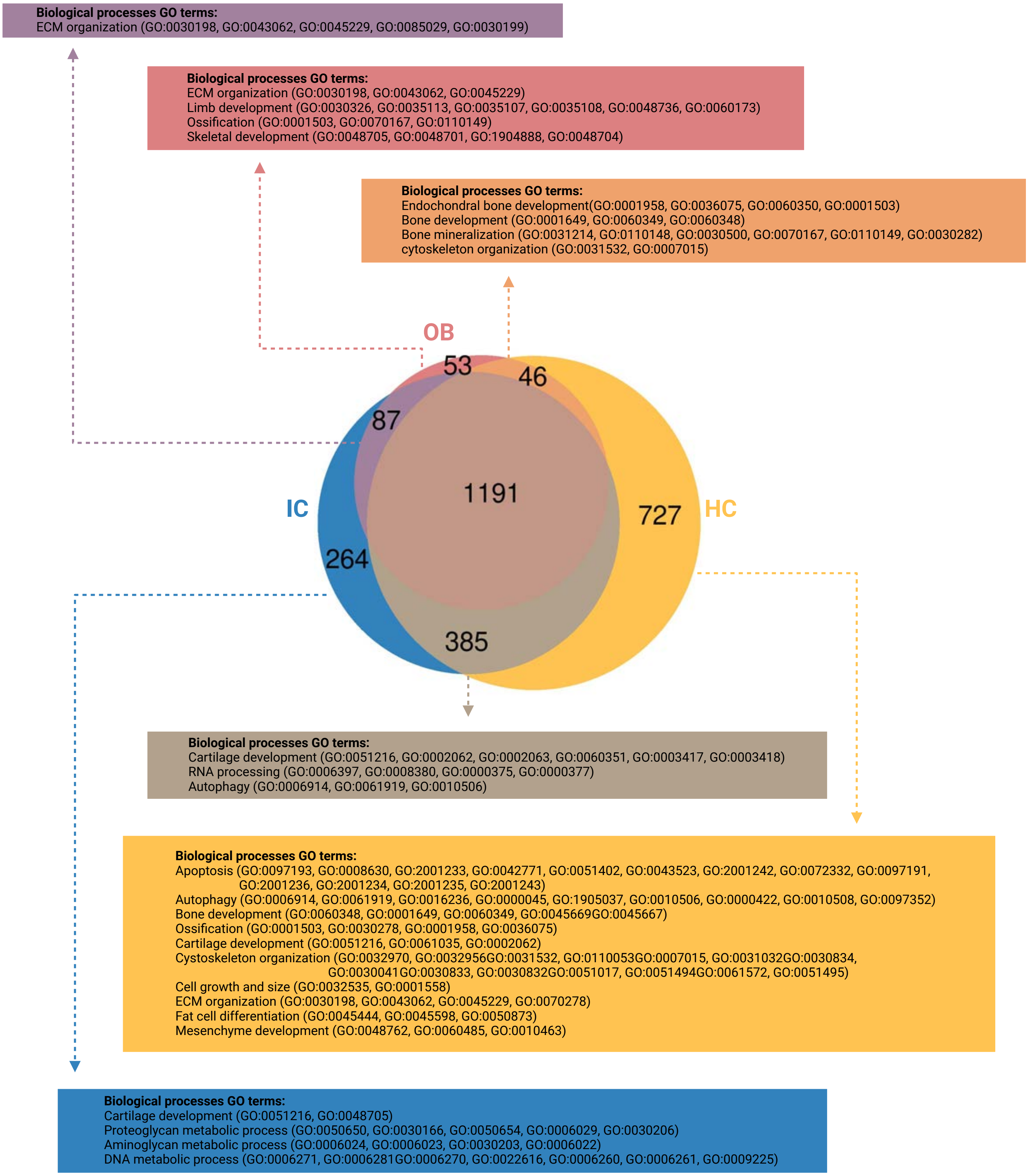


**Figure S6. GO enrichment analysis for biological processes of genes that are unique and shared among IC, OB and HC.** Venn diagram was taken from the last figure. All enrichment analyses were corrected by BH (p ≤ 0.05).

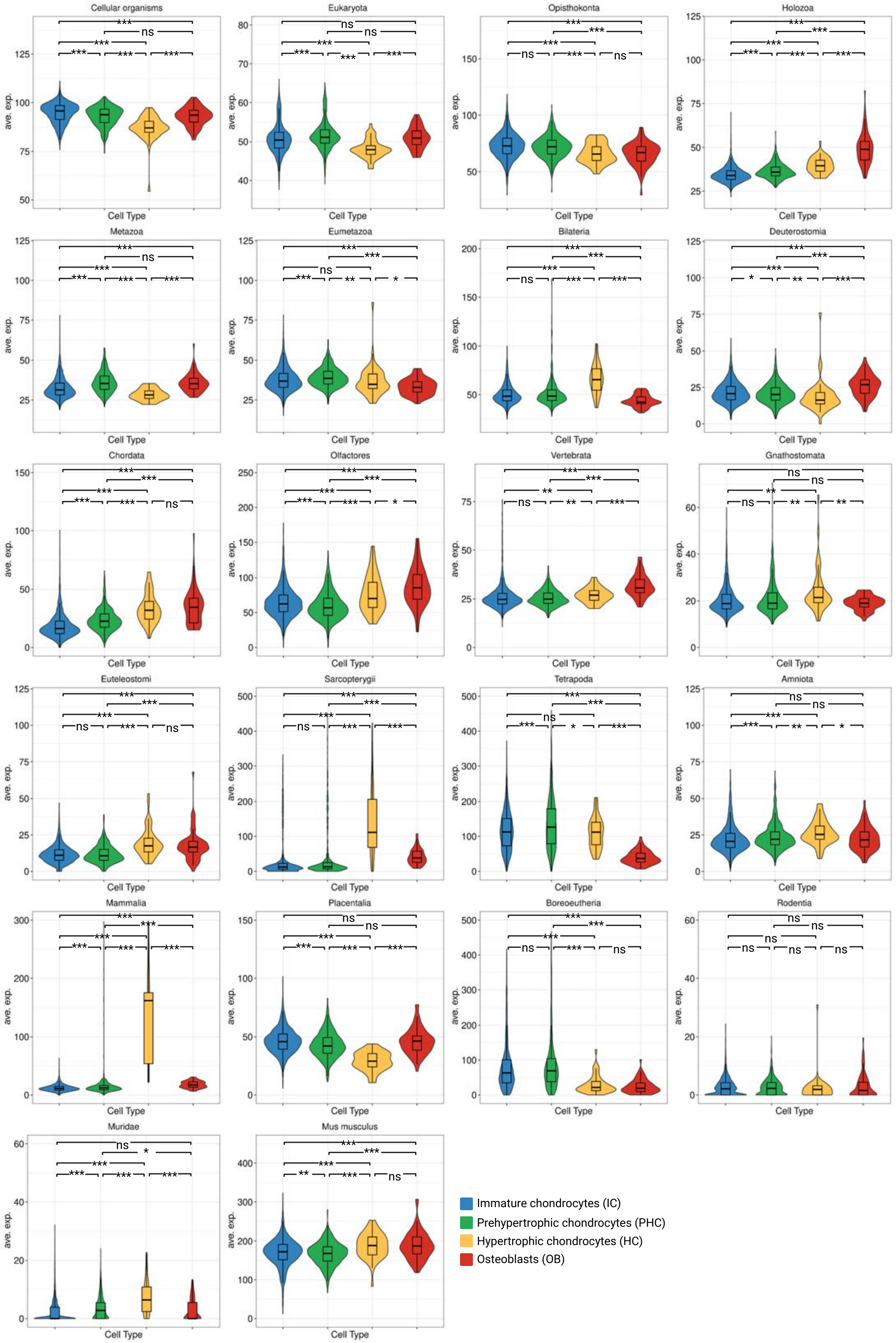

Cellular organisms
Eukaryota
Opisthokonta
Holozoa
Metazoa
Eumetazoa
Bilateria
Deuterostomia
Chordata
Olfactores
Vertebrata
Gnathostomata
Euteleostomi
Sarcopterygii
Tetrapoda
Amniota
Mammalia
Placentalia
Boreoeutheria
Rodentia
Muridae
Mus musculus
ave. exp.
Cell Type
Immature chondrocytes (IC)
Prehypertrophic chondrocytes (PHC)
Hypertrophic chondrocytes (HC)
Osteoblasts (OB)

**Figure S7. Average expression of genes in each ps in individual skeletal cell types.** The mean expression was calculated after grouping the genes expressed by each cell type according to their ps assignment. Statistical significance of average gene expression differences between the cell types in each PS was evaluated using a pairwise Wilcoxon test corrected for multiple comparisons by BH. Asterisks denote adjusted p value levels (* ≤0.05, **≤0.01, ***≤0.001).

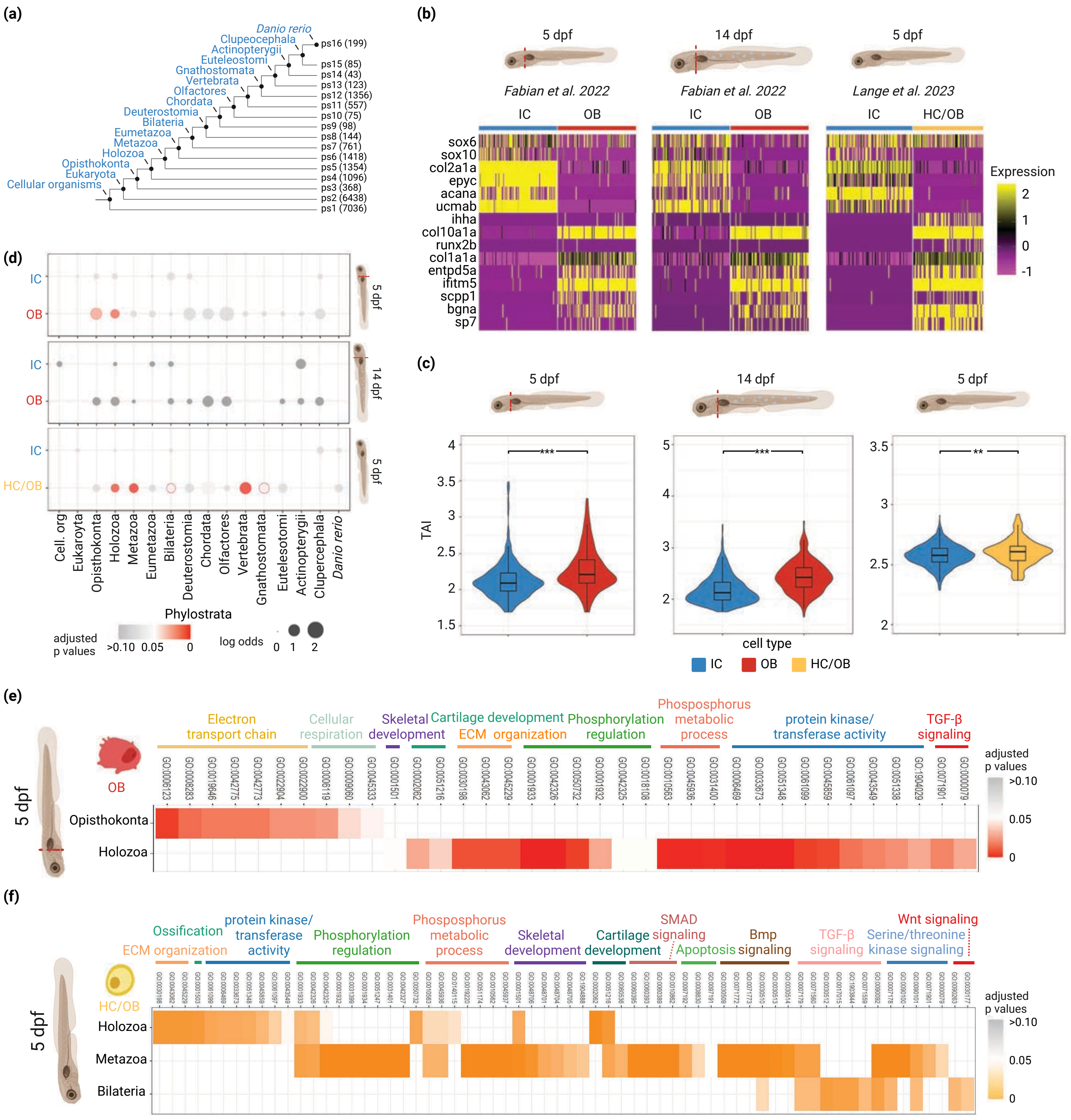

(a)
Danio rerio
Clupeocephala
Actinopterygii
Euteleostomi
Gnathostomata
Vertebrata
Olfactores
Chordata
Deuterostomia
Bilateria
Eumetazoa
Metazoa
Holozoa
Opisthokonta
Eukaryota
Cellular organisms
ps16 (199)
ps15 (85)
ps14 (43)
ps13 (123)
ps12 (1356)
ps11 (557)
ps10 (75)
ps9 (98)
ps8 (144)
ps7 (761)
ps6 (1418)
ps5 (1354)
ps4 (1096)
ps3 (368)
ps2 (6438)
ps1 (7036)
(b)
5 dpf
14 dpf
5 dpf
Fabian et al. 2022
Fabian et al. 2022
Lange et al. 2023
IC
OB
IC
OB
IC
HC/OB
sox6
sox10
col2a1a
epyc
acana
ucmab
ihha
col10a1a
runx2b
col1a1a
entpd5a
ifitm5
scpp1
bgna
sp7
Expression
2
1
0
-1
(c)
5 dpf
14 dpf
5 dpf
TAI
***
***
**
cell type
IC
OB
HC/OB
(d)
IC
OB
IC
OB
IC
HC/OB
5 dpf
14 dpf
5 dpf
Cell. org
Eukaroyta
Opisthokonta
Holozoa
Metazoa
Eumetazoa
Bilateria
Deuterostomia
Chordata
Olfactores
Vertebrata
Gnathostomata
Eutelesotomi
Actinopterygii
Clupercephala
Danio rerio
Phylostrata
adjusted p values
>0.10 0.05 0
log odds
0 1 2
(e)
5 dpf
OB
Electron transport chain
Cellular respiration
Skeletal development
Cartilage development
ECM organization
Phosphorylation regulation
Phosposphorus metabolic process
protein kinase/ transferase activity
TGF-β signaling
Opisthokonta
Holozoa
adjusted p values
>0.10
0.05
0
(f)
5 dpf
HC/OB
Ossification
ECM organization
protein kinase/ transferase activity
Phosphorylation regulation
Phosposphorus metabolic process
Skeletal development
Cartilage development
SMAD signaling
Apoptosis
Bmp signaling
TGF-β signaling
Serine/threonine kinase signaling
Wnt signaling
Holozoa
Metazoa
Bilateria
adjusted p values
>0.10
0.05
0

**Figure S8. Phylotranscriptomic analysis of skeletal cell types in zebrafish.** (a) Phylostratigraphy map of the zebrafish protein-coding genes. Each phylostratum (ps) number corresponds to a node in the phylogeny. Numbers in the parentheses indicate the number of genes that originated in the given phylostratum. (b) Gene expression profile of skeletal cell types using specific marker genes. The cell types were recovered from three single-cell transcriptome datasets sampled from 5dpf and 14dpf stages. (c) TAI profile of skeletal cell types in each dataset. IC, immature chondrocytes; OB, osteoblasts; HC, hypertrophic chondrocytes. The statistical significance of differences between TAI values was evaluated using a pairwise Wilcoxon test corrected for multiple comparisons by BH. Asterisks denote adjusted *p* value levels (* ≤0.05, **≤0.01, ***≤0.001). (d) Enrichment analyses of upregulated DEG distribution across the phylogeny. The representation of lineage-restricted genes associated with the evolution of a particular cell type is shown for each ps in log-odds values. Orange-to-red circles indicate statistically significant enrichment of genes associated with the evolution of a particular cell type, while grey circles indicate enrichments that are not statistically significant. Enrichments were tested using a two-tailed hypergeometric test corrected for multiple comparisons by FDR ($p \leq 0.05$). (e, f) GO enrichment analysis for biological processes of DEGs originated from the significantly enriched phylostrata in Fig. S8d. All enrichment analyses were corrected by BH ($p \leq 0.05$). Enriched phylostrata such as Vertebrata and Gnathostomata in HC/OB did not provide any significant GO terms.

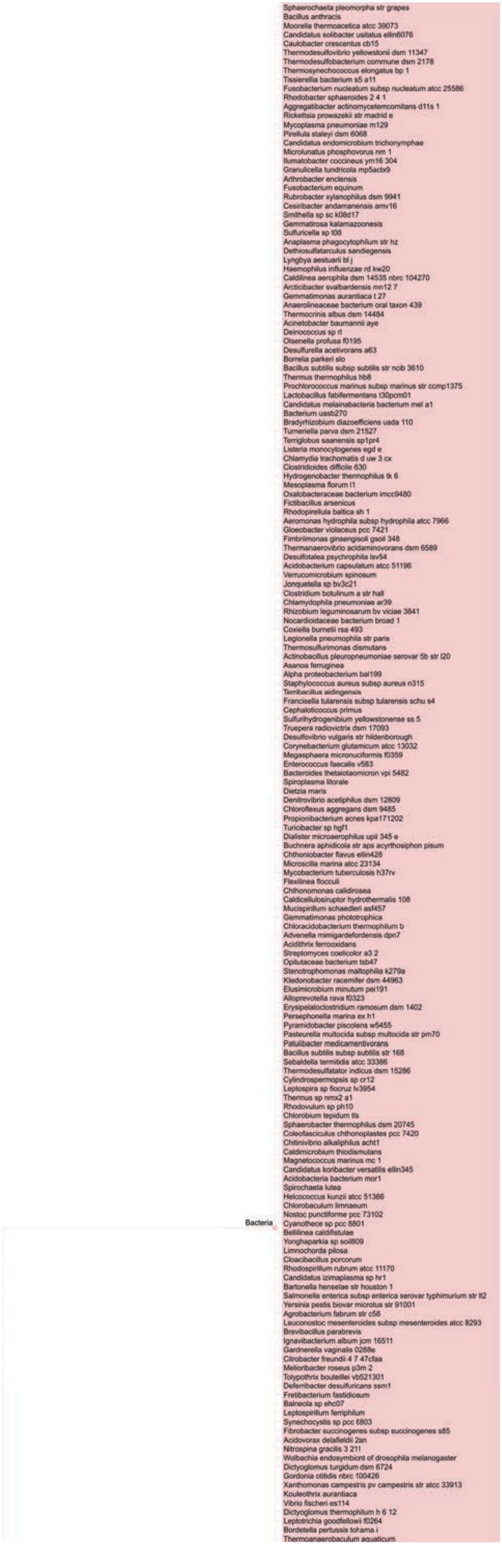
Bacteria
Sphaerochaeta pleomorpha str grapes
Bacillus anthracis
Moorella thermoacetica atcc 39073
Candidatus solibacter usitatus ellin6076
Caulobacter crescentus cb15
Thermodesulfovibrio yellowstonii dsm 11347
Thermodesulfobacterium commune dsm 2178
Thermosynechococcus elongatus bp 1
Tissierellia bacterium s5 a11
Fusobacterium nucleatum subsp nucleatum atcc 25586
Rhodobacter sphaeroides 2 4 1
Aggregatibacter actinomycetemcomitans d11s 1
Rickettsia prowazekii str madrid e
Mycoplasma pneumoniae m129
Pirellula staleyi dsm 6068
Candidatus endomicrobium trichonymphae
Microlunatus phosphovorus nm 1
Ilumatobacter coccineus ym16 304
Granulicella tundricola mp5actx9
Arthrobacter enclensis
Fusobacterium equinum
Rubrobacter xylanophilus dsm 9941
Cesiribacter andamanensis amv16
Smithella sp sc k08d17
Gemmatirosa kalamazoonesis
Sulfuricella sp t08
Anaplasma phagocytophilum str hz
Dethiosulfatarculus sandiegensis
Lyngbya aestuarii bl j
Haemophilus influenzae rd kw20
Caldilinea aerophila dsm 14535 nbrc 104270
Arcticibacter svalbardensis mn12 7
Gemmatimonas aurantiaca t 27
Anaerolineaceae bacterium oral taxon 439
Thermocrinis albus dsm 14484
Acinetobacter baumannii aye
Deinococcus sp rl
Olsenella profusa f0195
Desulfurella acetivorans a63
Borrelia parkeri slo
Bacillus subtilis subsp subtilis str ncib 3610
Thermus thermophilus hb8
Prochlorococcus marinus subsp marinus str ccmp1375
Lactobacillus fabifermentans t30pcm01
Candidatus melainabacteria bacterium mel a1
Bacterium uasb270
Bradyrhizobium diazoefficiens usda 110
Turneriella parva dsm 21527
Terriglobus saanensis sp1pr4
Listeria monocytogenes egd e
Chlamydia trachomatis d uw 3 cx
Clostridioides difficile 630
Hydrogenobacter thermophilus tk 6
Mesoplasma florum l1
Oxalobacteraceae bacterium imcc9480
Fictibacillus arsenicus
Rhodopirellula baltica sh 1
Aeromonas hydrophila subsp hydrophila atcc 7966
Gloeobacter violaceus pcc 7421
Fimbriimonas ginsengisoli gsoil 348
Thermanaerovibrio acidaminovorans dsm 6589
Desulfotalea psychrophila lsv54
Acidobacterium capsulatum atcc 51196
Verrucomicrobium spinosum
Jonquetella sp bv3c21
Clostridium botulinum a str hall
Chlamydophila pneumoniae ar39
Rhizobium leguminosarum bv viciae 3841
Nocardioidaceae bacterium broad 1
Coxiella burnetii rsa 493
Legionella pneumophila str paris
Thermosulfurimonas dismutans
Actinobacillus pleuropneumoniae serovar 5b str l20
Asanoa ferruginea
Alpha proteobacterium bal199
Staphylococcus aureus subsp aureus n315
Terribacillus aidingensis
Francisella tularensis subsp tularensis schu s4
Cephaloticoccus primus
Sulfurihydrogenibium yellowstonense ss 5
Truepera radiovictrix dsm 17093
Desulfovibrio vulgaris str hildenborough
Corynebacterium glutamicum atcc 13032
Megasphaera micronuciformis f0359
Enterococcus faecalis v583
Bacteroides thetaiotaomicron vpi 5482
Spiroplasma litorale
Dietzia maris
Denitrovibrio acetiphilus dsm 12809
Chloroflexus aggregans dsm 9485
Propionibacterium acnes kpa171202
Turicibacter sp hgf1
Dialister microaerophilus upii 345 e
Buchnera aphidicola str aps acyrthosiphon pisum
Chthoniobacter flavus ellin428
Microscilla marina atcc 23134
Mycobacterium tuberculosis h37rv
Flexilinea flocculi
Chthonomonas calidirosea
Caldicellulosiruptor hydrothermalis 108
Mucispirillum schaedleri asf457
Gemmatimonas phototrophica
Chloracidobacterium thermophilum b
Advenella mimigardefordensis dpn7
Acidithrix ferrooxidans
Streptomyces coelicolor a3 2
Opitutaceae bacterium tsb47
Stenotrophomonas maltophilia k279a
Ktedonobacter racemifer dsm 44963
Elusimicrobium minutum pei191
Alloprevotella rava f0323
Erysipelatoclostridium ramosum dsm 1402
Persephonella marina ex h1
Pyramidobacter piscolens w5455
Pasteurella multocida subsp multocida str pm70
Patulibacter medicamentivorans
Bacillus subtilis subsp subtilis str 168
Sebaldella termitidis atcc 33386
Thermodesulfatator indicus dsm 15286
Cylindrospermopsis sp cr12
Leptospira sp fiocruz lv3954
Thermus sp nmx2 a1
Rhodovulum sp ph10
Chlorobium tepidum tls
Sphaerobacter thermophilus dsm 20745
Coleofasciculus chthonoplastes pcc 7420
Chitinivibrio alkaliphilus acht1
Caldimicrobium thiodismutans
Magnetococcus marinus mc 1
Candidatus koribacter versatilis ellin345
Acidobacteria bacterium mor1
Spirochaeta lutea
Helcococcus kunzii atcc 51366
Chlorobaculum limnaeum
Nostoc punctiforme pcc 73102
Cyanothece sp pcc 8801
Bellilinea caldifistulae
Yonghaparkia sp soil809
Limnochorda pilosa
Cloacibacillus porcorum
Rhodospirillum rubrum atcc 11170
Candidatus izimaplasma sp hr1
Bartonella henselae str houston 1
Salmonella enterica subsp enterica serovar typhimurium str lt2
Yersinia pestis biovar microtus str 91001
Agrobacterium fabrum str c58
Leuconostoc mesenteroides subsp mesenteroides atcc 8293
Brevibacillus parabrevis
Ignavibacterium album jcm 16511
Gardnerella vaginalis 0288e
Citrobacter freundii 4 7 47cfaa
Melioribacter roseus p3m 2
Tolypothrix bouteillei vb521301
Deferribacter desulfuricans ssm1
Fretibacterium fastidiosum
Balneola sp ehc07
Leptospirillum ferriphilum
Synechocystis sp pcc 6803
Fibrobacter succinogenes subsp succinogenes s85
Acidovorax delafieldii 2an
Nitrospina gracilis 3 211
Wolbachia endosymbiont of drosophila melanogaster
Dictyoglomus turgidum dsm 6724
Gordonia otitidis nbrc 100426
Xanthomonas campestris pv campestris str atcc 33913
Kouleothrix aurantiaca
Vibrio fischeri es114
Dictyoglomus thermophilum h 6 12
Leptotrichia goodfellowii f0264
Bordetella pertussis tohama i
Thermoanaerobaculum aquaticum

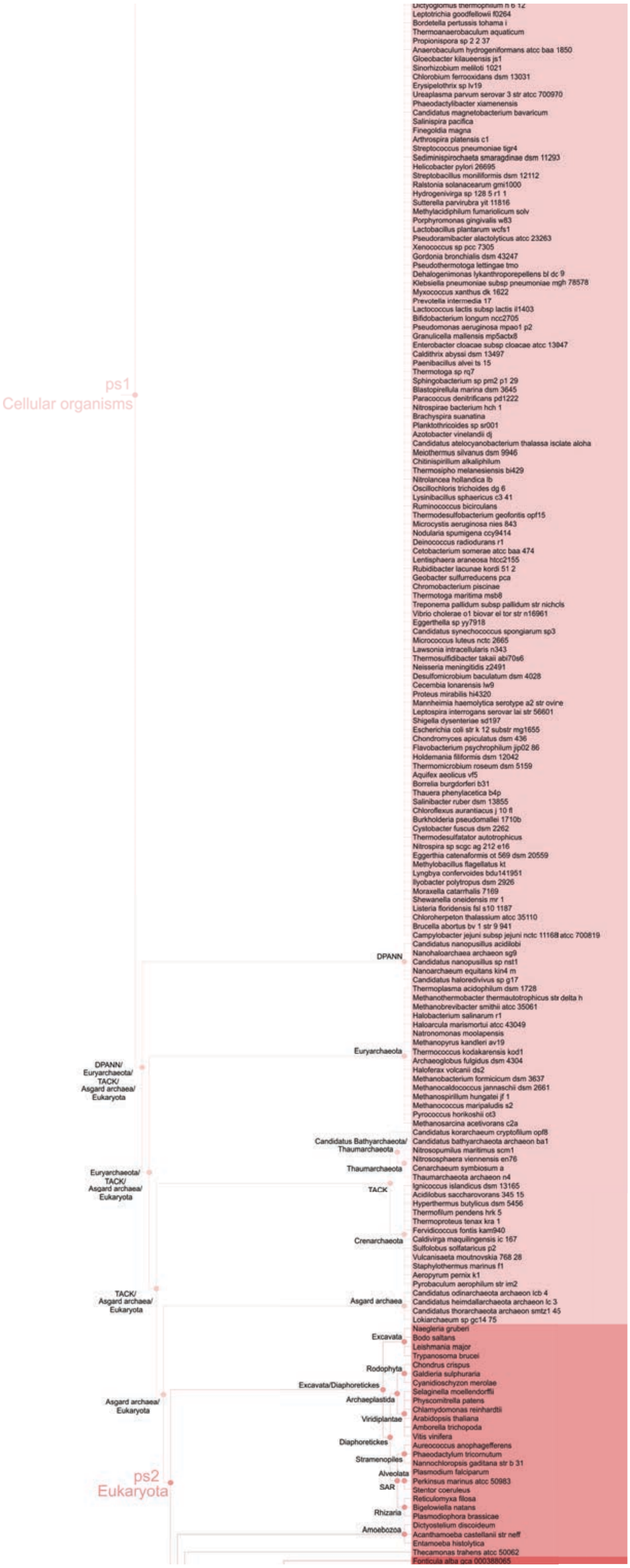

ps1
Cellular organisms
Dictyoglomus thermophilum h 6 12
Leptotrichia goodfellowii f0264
Bordetella pertussis tohama i
Thermoanaerobaculum aquaticum
Propionispora sp 2 2 37
Anaerobaculum hydrogeniformans atcc baa 1850
Gloeobacter kilaueensis js1
Sinorhizobium meliloti 1021
Chlorobium ferrooxidans dsm 13031
Erysipelothrix sp lv19
Ureaplasma parvum serovar 3 str atcc 700970
Phaeodactylibacter xiamenensis
Candidatus magnetobacterium bavaricum
Salinispira pacifica
Finegoldia magna
Arthrospira platensis c1
Streptococcus pneumoniae tigr4
Sediminispirochaeta smaragdinae dsm 11293
Helicobacter pylori 26695
Streptobacillus moniliformis dsm 12112
Ralstonia solanacearum gmi1000
Hydrogenivirga sp 128 5 r1 1
Sutterella parvirubra yit 11816
Methylacidiphilum fumariolicum solv
Porphyromonas gingivalis w83
Lactobacillus plantarum wcfs1
Pseudoramibacter alactolyticus atcc 23263
Xenococcus sp pcc 7305
Gordonia bronchialis dsm 43247
Pseudothermotoga lettingae tmo
Dehalogenimonas lykanthroporepellens bl dc 9
Klebsiella pneumoniae subsp pneumoniae mgh 78578
Myxococcus xanthus dk 1622
Prevotella intermedia 17
Lactococcus lactis subsp lactis il1403
Bifidobacterium longum ncc2705
Pseudomonas aeruginosa mpao1 p2
Granulicella mallensis mp5actx8
Enterobacter cloacae subsp cloacae atcc 13047
Caldithrix abyssi dsm 13497
Paenibacillus alvei ts 15
Thermotoga sp rq7
Sphingobacterium sp pm2 p1 29
Blastopirellula marina dsm 3645
Paracoccus denitrificans pd1222
Nitrospirae bacterium hch 1
Brachyspira suanatina
Planktothricoides sp sr001
Azotobacter vinelandii dj
Candidatus atelocyanobacterium thalassa isolate aloha
Meiothermus silvanus dsm 9946
Chitinispirillum alkaliphilum
Thermosipho melanesiensis bi429
Nitrolancea hollandica lb
Oscillochloris trichoides dg 6
Lysinibacillus sphaericus c3 41
Ruminococcus bicirculans
Thermodesulfobacterium geofontis opf15
Microcystis aeruginosa nies 843
Nodularia spumigena ccy9414
Deinococcus radiodurans r1
Cetobacterium somerae atcc baa 474
Lentisphaera araneosa htcc2155
Rubidibacter lacunae kordi 51 2
Geobacter sulfurreducens pca
Chromobacterium piscinae
Thermotoga maritima msb8
Treponema pallidum subsp pallidum str nichols
Vibrio cholerae o1 biovar el tor str n16961
Eggerthella sp yy7918
Candidatus synechococcus spongiarum sp3
Micrococcus luteus nctc 2665
Lawsonia intracellularis n343
Thermosulfidibacter takaii abi70s6
Neisseria meningitidis z2491
Desulfomicrobium baculatum dsm 4028
Cecembia lonarensis lw9
Proteus mirabilis hi4320
Mannheimia haemolytica serotype a2 str ovine
Leptospira interrogans serovar lai str 56601
Shigella dysenteriae sd197
Escherichia coli str k 12 substr mg1655
Chondromyces apiculatus dsm 436
Flavobacterium psychrophilum jip02 86
Holdemania filiformis dsm 12042
Thermomicrobium roseum dsm 5159
Aquifex aeolicus vf5
Borrelia burgdorferi b31
Thauera phenylacetica b4p
Salinibacter ruber dsm 13855
Chloroflexus aurantiacus j 10 fl
Burkholderia pseudomallei 1710b
Cystobacter fuscus dsm 2262
Thermodesulfatator autotrophicus
Nitrospira sp scgc ag 212 e16
Eggerthia catenaformis ot 569 dsm 20559
Methylobacillus flagellatus kt
Lyngbya confervoides bdu141951
Ilyobacter polytropus dsm 2926
Moraxella catarrhalis 7169
Shewanella oneidensis mr 1
Listeria floridensis fsl s10 1187
Chloroherpeton thalassium atcc 35110
Brucella abortus bv 1 str 9 941
Campylobacter jejuni subsp jejuni nctc 11168 atcc 700819
Candidatus nanopusillus acidilobi
DPANN
Nanohaloarchaea archaeon sg9
Candidatus nanopusillus sp nst1
Nanoarchaeum equitans kin4 m
Candidatus haloredivivus sp g17
Thermoplasma acidophilum dsm 1728
Methanothermobacter thermautotrophicus str delta h
Methanobrevibacter smithii atcc 35061
Halobacterium salinarum r1
Haloarcula marismortui atcc 43049
Natronomonas moolapensis
Methanopyrus kandleri av19
Euryarchaeota
Thermococcus kodakarensis kod1
Archaeoglobus fulgidus dsm 4304
Haloferax volcanii ds2
Methanobacterium formicicum dsm 3637
Methanocaldococcus jannaschii dsm 2661
Methanospirillum hungatei jf 1
Methanococcus maripaludis s2
Pyrococcus horikoshii ot3
Methanosarcina acetivorans c2a
DPANN/
Euryarchaeota/
TACK/
Asgard archaea/
Eukaryota
Candidatus korarchaeum cryptofilum opf8
Candidatus Bathyarchaeota/
Thaumarchaeota
Candidatus bathyarchaeota archaeon ba1
Nitrosopumilus maritimus scm1
Nitrososphaera viennensis en76
Thaumarchaeota
Cenarchaeum symbiosum a
Thaumarchaeota archaeon n4
Euryarchaeota/
TACK/
Asgard archaea/
Eukaryota
TACK
Ignicoccus islandicus dsm 13165
Acidilobus saccharovorans 345 15
Hyperthermus butylicus dsm 5456
Thermofilum pendens hrk 5
Thermoproteus tenax kra 1
Fervidicoccus fontis kam940
Crenarchaeota
Caldivirga maquilingensis ic 167
Sulfolobus solfataricus p2
Vulcanisaeta moutnovskia 768 28
Staphylothermus marinus f1
Aeropyrum pernix k1
Pyrobaculum aerophilum str im2
TACK/
Asgard archaea/
Eukaryota
Candidatus odinarchaeota archaeon lcb 4
Asgard archaea
Candidatus heimdallarchaeota archaeon lc 3
Candidatus thorarchaeota archaeon smtz1 45
Lokiarchaeum sp gc14 75
Naegleria gruberi
Excavata
Bodo saltans
Leishmania major
Trypanosoma brucei
Chondrus crispus
Rodophyta
Galdieria sulphuraria
Cyanidioschyzon merolae
Excavata/Diaphoretickes
Selaginella moellendorffii
Asgard archaea/
Eukaryota
Archaeplastida
Physcomitrella patens
Chlamydomonas reinhardtii
Viridiplantae
Arabidopsis thaliana
Amborella trichopoda
Vitis vinifera
Diaphoretickes
Aureococcus anophagefferens
Phaeodactylum tricornutum
Stramenopiles
Nannochloropsis gaditana str b 31
Alveolata
Plasmodium falciparum
ps2
Eukaryota
SAR
Perkinsus marinus atcc 50983
Stentor coeruleus
Reticulomyxa filosa
Bigelowiella natans
Rhizaria
Plasmodiophora brassicae
Dictyostelium discoideum
Amoebozoa
Acanthamoeba castellanii str neff
Entamoeba histolytica
Thecamonas trahens atcc 50062
Fonticula alba gca 000388065

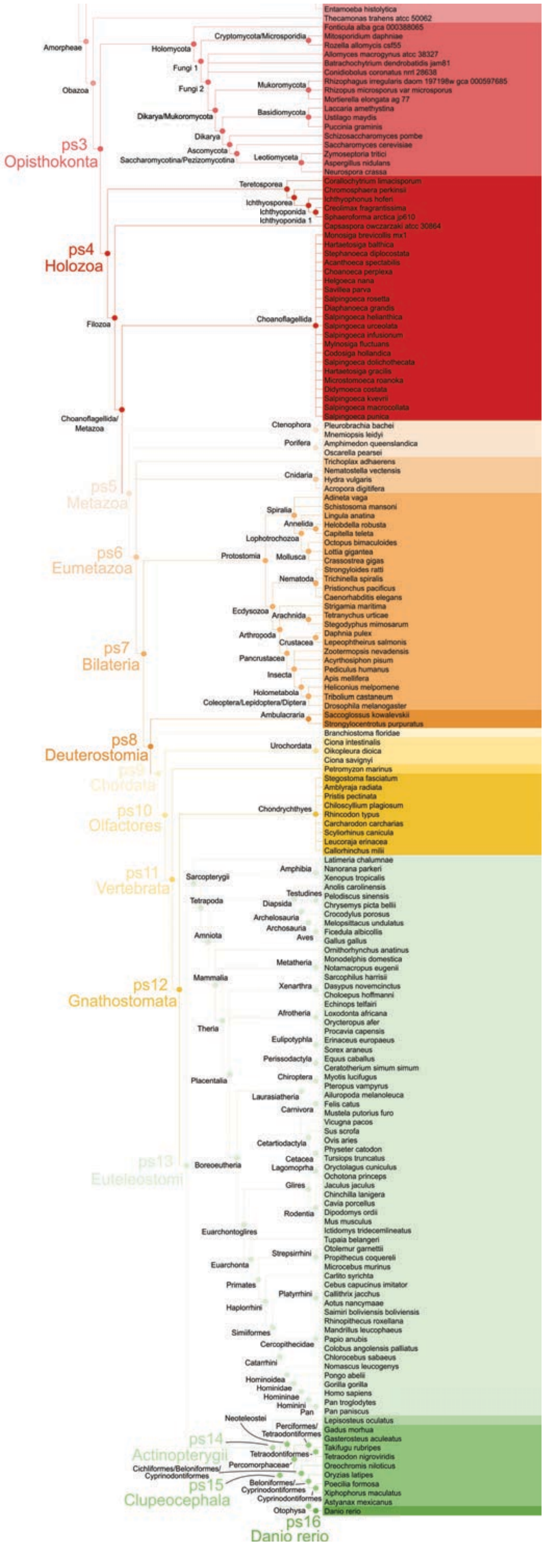

Amorpheae
Obazoa
Holomycota
Cryptomycota/Microsporidia
Fungi 1
Fungi 2
Mukoromycota
Dikarya/Mukoromycota
Basidiomycota
Dikarya
Ascomycota
Saccharomycotina/Pezizomycotina
Leotiomyceta
ps3
Opisthokonta
Teretosporea
Ichthyosporea
Ichthyoponida
Ichthyoponida 1
ps4
Holozoa
Filozoa
Choanoflagellida
Choanoflagellida/Metazoa
Ctenophora
Porifera
Cnidaria
ps5
Metazoa
Spiralia
Annelida
Lophotrochozoa
Protostomia
Mollusca
ps6
Eumetazoa
Nematoda
Ecdysozoa
Arachnida
Arthropoda
Crustacea
ps7
Bilateria
Pancrustacea
Insecta
Holometabola
Coleoptera/Lepidoptera/Diptera
Ambulacraria
ps8
Deuterostomia
Urochordata
ps9
Chordata
Chondrychthyes
ps10
Olfactores
ps11
Vertebrata
Sarcopterygii
Amphibia
Tetrapoda
Testudines
Diapsida
Archelosauria
Archosauria
Amniota
Aves
Metatheria
Mammalia
Xenarthra
ps12
Gnathostomata
Afrotheria
Theria
Eulipotyphla
Perissodactyla
Placentalia
Chiroptera
Laurasiatheria
Carnivora
Cetartiodactyla
ps13
Euteleostomi
Boreoeutheria
Cetacea
Lagomorpha
Glires
Rodentia
Euarchontoglires
Strepsirrhini
Euarchonta
Primates
Platyrrhini
Haplorrhini
Simiiformes
Cercopithecidae
Catarrhini
Hominoidea
Hominidae
Homininae
Hominini
Pan
Neoteleostei
Perciformes/Tetraodontiformes
ps14
Actinopterygii
Tetraodontiformes
Cichliformes/Beloniformes/Cyprinodontiformes
Percomorphaceae
Beloniformes/Cyprinodontiformes
ps15
Clupeocephala
Cyprinodontiformes
Otophysa
ps16
Danio rerio
Entamoeba histolytica
Thecamonas trahens atcc 50062
Fonticula alba gca 000388065
Mitosporidium daphniae
Rozella allomycis csf55
Allomyces macrogynus atcc 38327
Batrachochytrium dendrobatidis jam81
Conidiobolus coronatus nrrl 28638
Rhizophagus irregularis daom 197198w gca 000597685
Rhizopus microsporus var microsporus
Mortierella elongata ag 77
Laccaria amethystina
Ustilago maydis
Puccinia graminis
Schizosaccharomyces pombe
Saccharomyces cerevisiae
Zymoseptoria tritici
Aspergillus nidulans
Neurospora crassa
Corallochytrium limacisporum
Chromosphaera perkinsii
Ichthyophonus hoferi
Creolimax fragrantissima
Sphaeroforma arctica jp610
Capsaspora owczarzaki atcc 30864
Monosiga brevicollis mx1
Hartaetosiga balthica
Stephanoeca diplocostata
Acanthoeca spectabilis
Choanoeca perplexa
Helgoeca nana
Savillea parva
Salpingoeca rosetta
Diaphanoeca grandis
Salpingoeca helianthica
Salpingoeca urceolata
Salpingoeca infusionum
Mylnosiga fluctuans
Codosiga hollandica
Salpingoeca dolichothecata
Hartaetosiga gracilis
Microstomoeca roanoka
Didymoeca costata
Salpingoeca kvevrii
Salpingoeca macrocollata
Salpingoeca punica
Pleurobrachia bachei
Mnemiopsis leidyi
Amphimedon queenslandica
Oscarella pearsei
Trichoplax adhaerens
Nematostella vectensis
Hydra vulgaris
Acropora digitifera
Adineta vaga
Schistosoma mansoni
Lingula anatina
Helobdella robusta
Capitella teleta
Octopus bimaculoides
Lottia gigantea
Crassostrea gigas
Strongyloides ratti
Trichinella spiralis
Pristionchus pacificus
Caenorhabditis elegans
Strigamia maritima
Tetranychus urticae
Stegodyphus mimosarum
Daphnia pulex
Lepeophtheirus salmonis
Zootermopsis nevadensis
Acyrthosiphon pisum
Pediculus humanus
Apis mellifera
Heliconius melpomene
Tribolium castaneum
Drosophila melanogaster
Saccoglossus kowalevskii
Strongylocentrotus purpuratus
Branchiostoma floridae
Ciona intestinalis
Oikopleura dioica
Ciona savignyi
Petromyzon marinus
Stegostoma fasciatum
Amblyraja radiata
Pristis pectinata
Chiloscyllium plagiosum
Rhincodon typus
Carcharodon carcharias
Scyliorhinus canicula
Leucoraja erinacea
Callorhinchus milii
Latimeria chalumnae
Nanorana parkeri
Xenopus tropicalis
Anolis carolinensis
Pelodiscus sinensis
Chrysemys picta bellii
Crocodylus porosus
Melopsittacus undulatus
Ficedula albicollis
Gallus gallus
Ornithorhynchus anatinus
Monodelphis domestica
Notamacropus eugenii
Sarcophilus harrisii
Dasypus novemcinctus
Choloepus hoffmanni
Echinops telfairi
Loxodonta africana
Orycteropus afer
Procavia capensis
Erinaceus europaeus
Sorex araneus
Equus caballus
Ceratotherium simum simum
Myotis lucifugus
Pteropus vampyrus
Ailuropoda melanoleuca
Felis catus
Mustela putorius furo
Vicugna pacos
Sus scrofa
Ovis aries
Physeter catodon
Tursiops truncatus
Oryctolagus cuniculus
Ochotona princeps
Jaculus jaculus
Chinchilla lanigera
Cavia porcellus
Dipodomys ordii
Mus musculus
Ictidomys tridecemlineatus
Tupaia belangeri
Otolemur garnettii
Propithecus coquereli
Microcebus murinus
Carlito syrichta
Cebus capucinus imitator
Callithrix jacchus
Aotus nancymaae
Saimiri boliviensis boliviensis
Rhinopithecus roxellana
Mandrillus leucophaeus
Papio anubis
Colobus angolensis palliatus
Chlorocebus sabaeus
Nomascus leucogenys
Pongo abelii
Gorilla gorilla
Homo sapiens
Pan troglodytes
Pan paniscus
Lepisosteus oculatus
Gadus morhua
Gasterosteus aculeatus
Takifugu rubripes
Tetraodon nigroviridis
Oreochromis niloticus
Oryzias latipes
Poecilia formosa
Xiphophorus maculatus
Astyanax mexicanus
Danio rerio

**Figure S9. Expanded consensus phylogeny used in the genomic phylostratigraphy analysis.** The tree covers divergence from the last common ancestor to zebrafish, *Danio rerio*. Sixteen nodes (phylostrata, ps) were considered in the analysis.
